# Hierarchy of prediction errors shapes the learning of context-dependent sensory representations

**DOI:** 10.1101/2024.09.30.615819

**Authors:** Matthias C. Tsai, Jasper Teutsch, Willem A.M. Wybo, Fritjof Helmchen, Abhishek Banerjee, Walter Senn

**Affiliations:** Department of Physiology, University of Bern, Bern, Switzerland; Brain Research Institute, University of Zürich, Zürich, Switzerland; Department of Pharmacology, University of Oxford, Oxford, UK; Blizard Institute, Queen Mary University of London, London, UK; Peter Grünberg Institute, Jülich Research Center, Jülich, Germany; Neuroscience Center Zürich, University of Zürich, Zürich, Switzerland; University Research Priority Program (URPP), Adaptive Brain Circuits in Development and Learning (AdaBD), University of Zürich, Zürich, Switzerland

**Keywords:** top-down modulation, representation learning, lateral orbitofrontal cortex, reversal learning, apical amplification, primary somatosensory cortex, continual learning, non-stationary environment, interneurons, disinhibition, online machine learning

## Abstract

How sensory information is interpreted depends on context, yet the neural mechanisms by which context shapes sensory processing remain poorly understood. To address this question, we developed a computational model constrained by *in vivo* functional imaging of cortical neurons in mice during reversal learning of a tactile sensory discrimination task. During learning, layer 2/3 somatosensory neurons enhanced their response to reward-predictive stimuli. The model accounted for these observations through selective top-down gain amplification of apical dendritic inputs, accompanied by reduced reward-prediction errors and increased confidence in outcome predictions. Upon rule-reversal, the lateral orbitofrontal cortex, through disinhibitory VIP interneurons, encoded a context-prediction error signaling a loss of confidence. The hierarchy of reward- and context-prediction errors across cortical areas is mirrored in top-down signals modulating apical activity of simulated pyramidal neurons in the primary sensory cortex. The model explains how contextual changes are detected and how reward- and context-prediction error signals, originating in different cortical regions, interact to reshape the sensory representation.

---

The theory of reward-prediction errors with their neuronal correlates represents a fundamental milestone for behavioral neuroscience [1]. It forms the foundation for reinforcement learning in the brain and inspired developments in artificial intelligence [2, 3]. Making reward predictions in complex environments, however, also requires the identification of ongoing contexts, i.e., unobservable latent variables whose correct inference is necessary for optimal behavior. For one context, the rewards in an environment might follow certain rules; in another context these rules might be altered, hence altering the reward contingency. Yet, it is not clear – from a normative point of view – how context is represented, how a change in context is even detected, and how a putative context change can reshape computation. Although the ultimate goal of reward-based learning lies in minimizing reward-prediction errors and selecting actions to maximize reward, computing context-prediction errors and reducing them constitutes an essential intermediary goal. Therefore, elucidating how reward- and context-prediction errors jointly modulate sensory processing is crucial to understand the neural mechanisms underlying adaptive behavior in the brain [4].

Being able to extract a large number of features from the sensory world is advantageous to make accurate decisions in a broad range of contexts. However, having a rich sensory representation also enlarges the search space from which to identify behaviorally relevant information. The brain addresses this dilemma by selectively amplifying task-relevant sensory responses [5–7] (Fig. 1a). While the origin of these enhanced responses in the sensory cortex is not fully understood, they can be modeled as multiplicative gain amplification [8]. Sensory response amplification depends on context, which indicates a top-down attentional modulation rather than a bottom-up adaptation [9–11]. Apical dendrites of pyramidal neurons receive top-down afferents [12], multiplicatively modulate the firing rate [13, 14] and acquire strengthened responses to stimuli that predict upcoming salient outcomes [15, 16]. Therefore, apical dendrites are suspected to selectively amplify neural responses in order to increase the internal salience of task-relevant sensory stimuli [17, 18].

The amplification of a select subset of features, however, can also have drawbacks. A once advantageous salience allocation can become inadequate after a change in context. Therefore, detecting such contextual changes becomes crucial for the adaptability of behavior. In animals, such scenarios are studied by means of reversal learning tasks [19]. Most prominently, lateral orbitofrontal cortex (lOFC) has been shown to respond to changes in reward contingency and plays a critical role in flexibly adjusting behavior [19, 20, 53]. Further work has shown that the lOFC can inhibit apical dendrites of pyramidal neurons in the primary visual (V1) and auditory cortex (A1) via somatostatin-expressing (SST) interneurons [21, 22], which could be a potential mechanism to revert acquired apical amplification patterns that are no longer appropriate after a change in context.

**Fig. 1.**
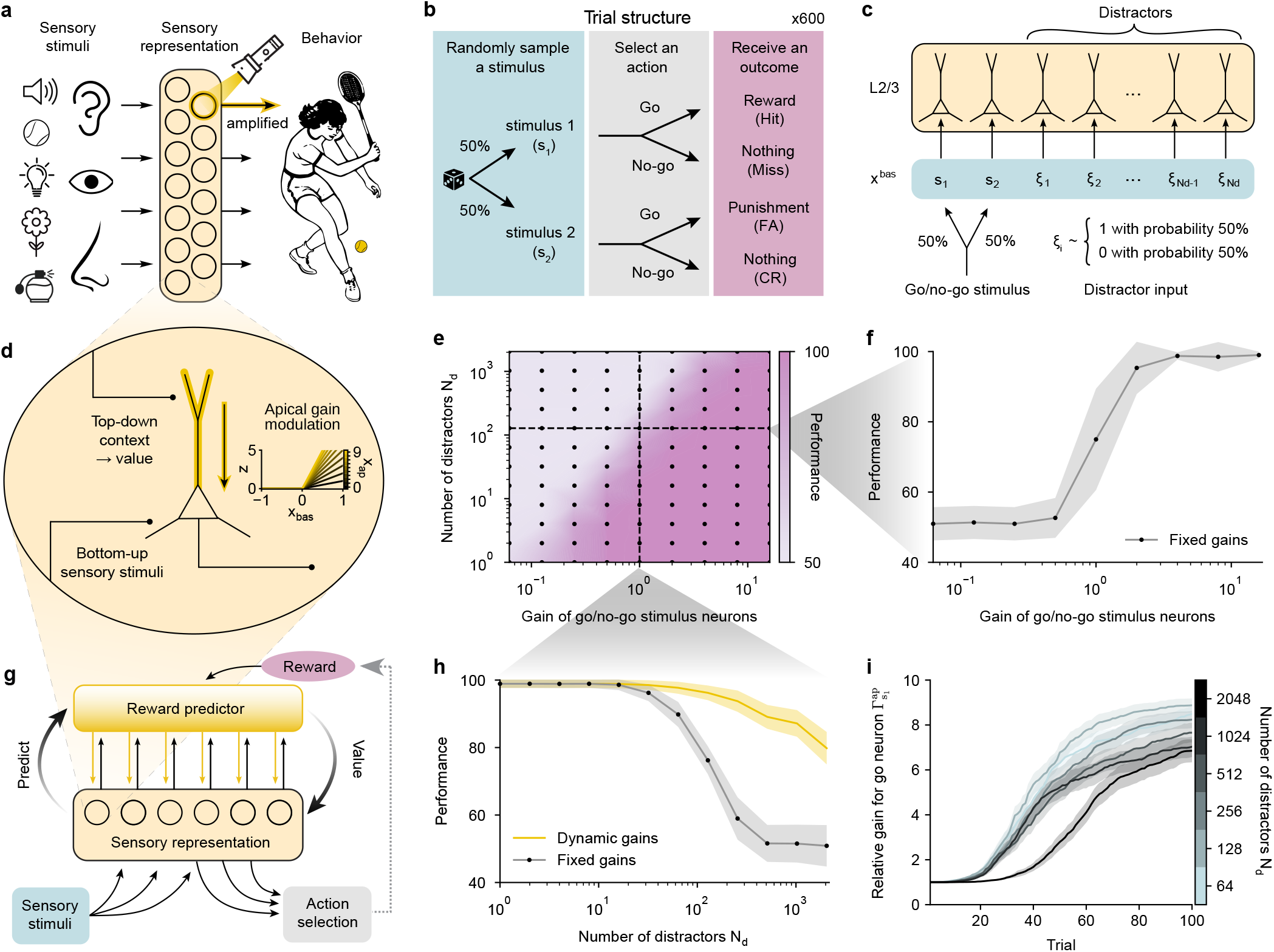
Value-based apical gain modulation amplifies task-relevant sensory signals. **a**, Sensory representations contain information about many sensory inputs and selectively amplify features that are most important for behavioral decisions. **b**, Structure of the go/no-go sensory discrimination task with each combination of stimulus and action leading to an outcome (Hit, receive reward; CR, correct rejection, omission of punishment; FA, false alarm, receive punishment; Miss, omission of reward). **c**, Bottom-up inputs to the sensory pyramidal neurons. The go (s_1_) and no-go (s_2_) stimuli drive separate neurons. Task-irrelevant features drive distractor neurons. The task-irrelevant distractor inputs are each modeled as Bernoulli random variables. **d**, Schematic showing how top-down afferents onto pyramidal neurons multiplicatively modulate the gain of the bottom-up input. In the model, the apical input reflects the neuron-specific value and the bottom-up input has a fixed selectivity. The plot shows the multiplicative effect of different apical dendrite activations *x*_ap_ on the firing rate *z* induced by the bottom-up activation *x*_bas_, see equation (1). **e**, Performance of the online policy gradient algorithm without dynamic gain adaptation on a go/no-go sensory discrimination task depending on the number of distractor neurons and on the fixed gain of the two sensory neurons selective for either s_1_ or s_2_. The gain of the distractor neurons was always set to 1. Performance was evaluated after 600 trials of training. Each black dot corresponds to the average from 64 different pseudorandom initializations. **f**, Horizontal slice of panel e with 128 distractor neurons and different fixed gains for the go and no-go stimulus encoding neurons. The gray area shows the standard deviation. **g**, Computational model of sensory processing. Bottom-up sensory stimuli drive the pyramidal neurons of the sensory representation. The bottom-up activation is amplified by top-down apical afferents conveying neuron-specific value. The values, derived from a reward prediction network, measure how much the sensory neuron activities are associated with future rewards. The sensory pyramidal neurons project both to the reward prediction network and to a policy network that learns to select actions through policy gradient. **h**, Performance achieved with and without dynamic gain modulation, depending on the number of distractor neurons. The values without gain adaptation correspond to the vertical slice in panel e. **i**, As learning progressed, the apical gain of the go-stimulus (s_1_) selective neuron increased, relative to the average gain of the other neurons, see equation (12). The mean and the standard error of this relative gain 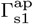 are shown for simulations with different numbers of distractors across the first 100 trials. With greater numbers of distractor neurons, the difficulty to identify reward-associated neurons increases, thus 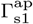 requires more trials to rise.

Here, we measured functional responses from the primary somatosensory cortex (S1) and lOFC neurons of mice during reversal learning. We integrated these findings into a computational model explaining how the lOFC can support behavioral flexibility by actively reshaping sensory representations. We examined the robustness of online reinforcement learning in the presence of a large set of irrelevant stimuli and show that the reward-guided apical gain amplification of sensory pyramidal neurons enhanced performance. Using this computational framework, we show that calcium signals in lOFC during stimulus presentation reflect a context-prediction error, which is enhanced upon rule reversal as a consequence of observing highly unexpected outcomes. S1 imaging during optogenetic lOFC inhibition indicates that the context-prediction error encoded by lOFC supports the adaptation to the new sensorimotor association rule. Using computational modeling supported by experimental evidence, we present two mechanisms by which the lOFC can reshape the top-down amplification of relevant sensory neurons learned during the previous rule. Finally, we demonstrate that silencing vasoactive intestinal polypeptide-expressing (VIP) interneurons in the lOFC impairs rule-switching. Taken together, our results show how the brain builds context-dependent sensory representations to support behavioral adaptation in non-stationary environments, and provide new perspectives on the roles of distinct inhibitory mechanisms in that process.

## Results

### Value-based apical amplification of sensory representations

Learning from large sensory representations containing few task-relevant and many task-irrelevant neurons constitutes a needle in the haystack problem. The task-irrelevant neurons act as distractors and obfuscate which signals to base decisions on. This applies at a high level, e.g., animals perceiving with multiple senses to navigate complex environments. However the issue grows considerably as the brain extracts diverse features from each sensory stimulus (Fig. 1a). We replicated this problem by simulating a rewarded go/no-go sensory discrimination task (see Methods, Simulated go/no-go task), where action selection was learned with online policy gradient (for a policy based on expected rewards see Extended Data Fig. S1, and Supplementary Methods 4). The occurrence of a stimulus s_1_ could lead to a reward if a specific action (go) was performed, and another stimulus s_2_ could lead to a punishment that could be avoided by not performing that go-action (Fig. 1b). The sensory representation consisted of rate-based two-compartment neurons. Two neurons each responded to a stimulus (s_1_ or s_2_) and varying numbers of distractor neurons were randomly activated (Fig. 1c). The activity in the apical compartment multiplicatively modulated the firing rate (Fig. 1d, see Gain modulation in sensory representations). To benchmark how gain modulation in the sensory representation could facilitate online policy learning, the gain of the s_1_ and s_2_ selective neurons was fixed to a range of values, while the gains of the distractor neurons remained fixed to 1 (see Supplementary Methods 3). For equal gains, the task was learned consistently in the presence of few distractors, but the learning performance began to falter around 100 distractor neurons (Fig. 1e). Differences in gain of the task-relevant neurons compared to the distractor neurons also affected learning. Weak task-relevant signals could spoil the performance, but the performance degradation caused by an overwhelming number of distractors could be rescued by increasing the gain of the s_1_ and s_2_ stimuli encoding neurons (Fig. 1f). In summary, the simulations demonstrate how improving sensory representations by amplifying the response of task-relevant sensory neurons supports behavioral learning.

In the brain, the amplification of task-relevant sensory responses has been reported to correlate with behavioral learning [6, 23]. Further experimental findings have shown that apical activity increases somatic excitability [13, 24], that learning enhances the apical response to behaviorally relevant stimuli [15, 16], and that adaptation in apical dendrites of sensory neurons precedes behavioral learning [25]. We therefore postulated that changes in sensory responses promoting task-relevant information could be modeled as top-down excitation of apical dendrites that multiplicatively amplifies bottom-up basal activations (Fig. 1d and Extended Data Fig. S2a). The guiding principles underlying this conditioning of the sensory representation remain unclear, but the enhancement of responses to stimuli that are predictive of future rewards has been reported with high consistency [19, 23, 26–28] (Extended Data Fig. S3a). We replicated these experimental observations by modeling pyramidal neurons that dynamically modulate bottom-up sensory inputs with top-down signals encoding the stimulus-reward association (Fig. 1d). A reward prediction network derives neuron-specific values that measure whether bottom-up inputs are associated with future rewards (Fig. 1g, see Apical amplification based on neuron-specific value). Sensory neurons are then amplified by shaping apical excitation according to this neuron-specific value, which encodes the expected relevance of each neuron for predicting future rewards (see Supplementary Equations 2). The simulated learning performances confirm that reward-guided apical gain modulation makes the system more robust to the presence of task-irrelevant stimuli (Fig. 1h). The resulting apical salience map increases the responses of sensory neurons according to their reward-association (Fig. 1i), and this in turn improves the sensory representation for action selection (see Extended Data Fig. S4 for a comparison with alternative algorithms). Hence, top-down projections onto apical dendrites of sensory neurons support behavioral learning by amplifying task-relevant information and increasing the signal-to-noise ratio (Extended Data Fig. S5).

### Lateral OFC signals a context-prediction error at stimulus time

So far, the model made use of reward predictions in a fixed stationary context. Next, we show how tracking the second-order statistics of reward-prediction errors is helpful in non-stationary environments, e.g., in case the rule of the go/no-go task is reversed (Fig. 2a). When exposed to a new task, animals may be agnostic about future outcomes, and any outcome can be considered equally uncertain. However, as an internal model of the task is constructed and the outcome predictions become more and more accurate, the uncertainty about the expected outcomes decrease. At the same time, any deviation from the predicted outcome becomes increasingly unexpected. How expected or unexpected an outcome is, can be captured by using a dynamic estimate 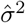 of the expected squared reward-prediction error *σ*^2^ (see Estimating the confidence of the reward prediction). The estimate 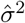 evaluates the variance unexplained by the model (see Supplementary Equations 4) and can be interpreted as the *prediction uncertainty* —quantifying the expected uncertainty of the reward predictions (see Supplementary Equations 3). As learning progresses and the reward predictions 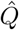 improve, 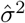 will decrease (Fig. 2b). In consequence, a reward after the go-stimulus (s_1_) will be predicted with increasing confidence, while any other outcome becomes less and less expected.

**Fig. 2.**
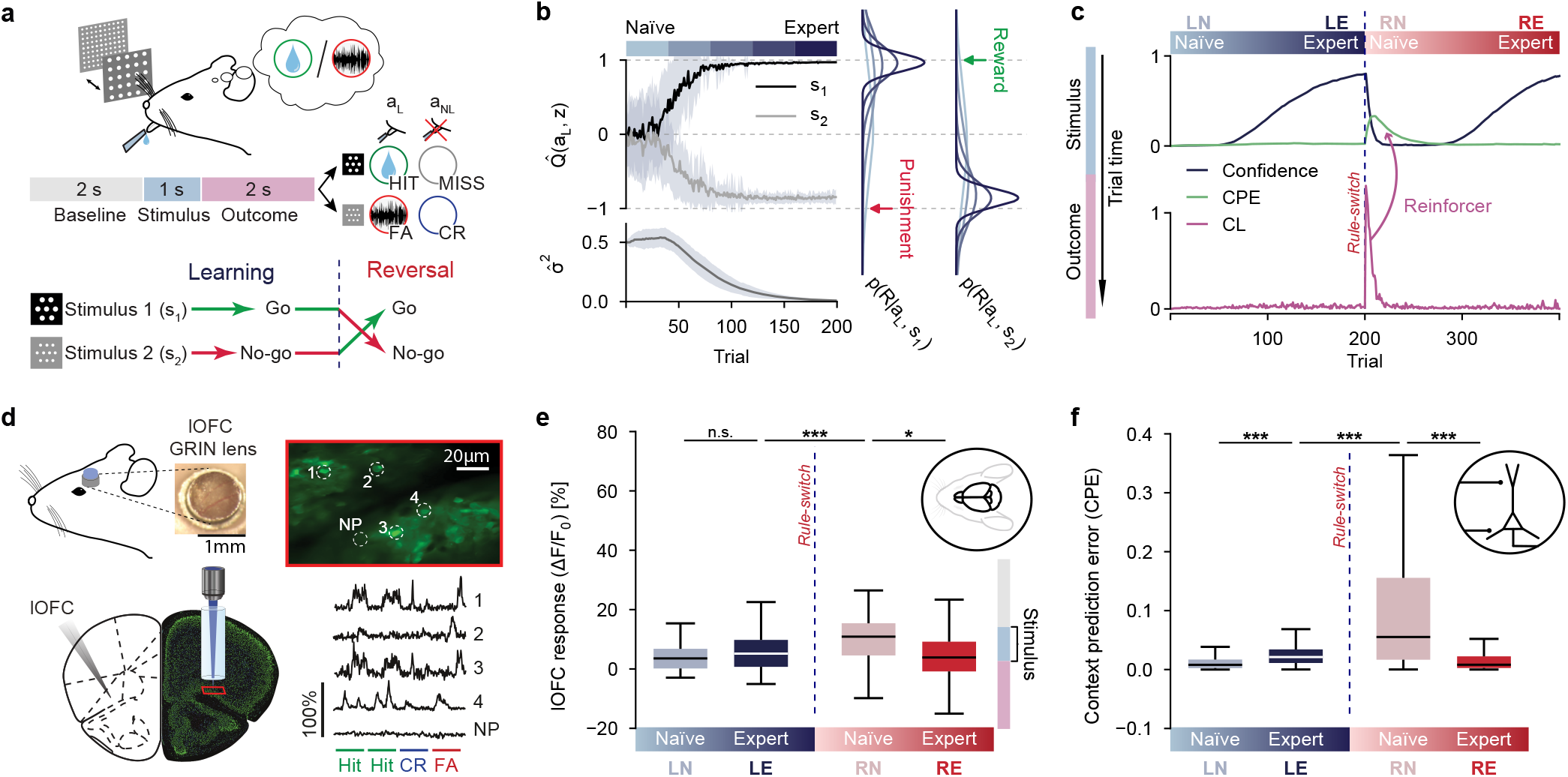
Lateral orbitofrontal cortex signals a context-prediction error at stimulus time. **a**, Go/no-go sensory discrimination task with rule reversal. Top: schematic of experimental setup, trial structure and possible outcome types (Hit, CR, FA, Miss), depending on a lick (*a*_L_) or no-lick (*a*_NL_). Bottom: after animals show stable expert discriminatory performance (learning), stimulus-outcome contingencies are reversed (reversal). **b**, Evolution of the simulated reward prediction 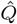 depending on the sensory representation *z* and the lick-action *a*_*L*_. Top: the values 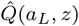 are shown across trials for the rewarded go-stimulus (s_1_) and for the punished no-go-stimulus (s_2_). Bottom: The estimate of the squared reward-prediction error 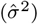 tracks the uncertainty of the reward prediction. Right: Estimated probability densities for the predicted reward as Gaussian distributions parameterized by the mean 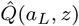 and variance 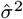 for either stimulus (s_1_ or s_2_). As learning progresses from the naïve (light blue) to the expert stage (dark blue), the estimated densities become more accurate and narrow. Colored arrows indicate unexpected outcomes in case of rule reversal, such as punishment following s_1_ (red) or reward following s_2_ (green). Details of the model an be found in the Methods and Supplementary Methods 3. **c**, Top: Prediction confidence (*c*, black) and context-prediction error (CPE, green, see equations 7-9) during the stimulus window, which occurs before the outcome window in trial time (see vertical time axis on the left). Bottom: The confidence loss (CL) triggered by unexpected outcomes. The CL entrains the CPE to appear during the stimulus window (purple ‘Reinforcer’ arrow pointing backwards in trial time, i.e. from the outcome to the stimulus window). As a result, few trials after the rule reversal, the CL ‘travels back’ within trial time and becomes a prospective CPE. **d**, Left: schematic and photograph of LOFC cannula and GRIN lens implantation. Right, two-photon fluorescence image of L2/3 lOFC neurons imaged through the GRIN lens and extracted traces of example cells and neuropil (NP). Traces of four concatenated example trials (whole trial) and corresponding trial outcome from an expert animal (LE). **e**, Box plots of Ca^2+^ transient amplitudes (ΔF*/*F_0_) from L2/3 lOFC neurons in the stimulus window during different learning phases of the reversal learning task (39, 122, 166, 170 neurons from 3 mice during the LN, LE, RN, RE phase respectively). The phases correspond to learning naïve (LN), learning expert (LE), reversal naïve (RN) and reversal expert (RE). Statistical differences were tested by two-sided Welch’s t-tests with Bonferroni correction, LN vs. LE (t(56)=-1.4, p_adj_=0.50; Δ=-1.7, 95% CI [-4.2, 0.75]), LE vs. RN (t(240)=-4.7, p_adj_*<*0.001; Δ=-4.1, 95% CI [-5.9, -2.4]), RN vs RE (t(241)=2.9, p_adj_=0.011; Δ=4.7, 95% CI [1.5, 7.8]). The asterisks indicate whether the p-values were ≥ 0.05 (n.s), *<*0.05 (*), *<*0.01 (**) or *<*0.001 (***). This convention is maintained throughout the manuscript. **f**, Same as in panel e, for the simulated context-prediction error; LN vs. LE (t(11604)=-45, p_adj_*<*0.001; Δ=-0.013, 95% CI [-0.014, -0.012]), LE vs. RN (t(6778)=-55, p_adj_*<*0.001; Δ=-0.077, 95% CI [-0.079, -0.074]), RN vs RE (t(6957)=61, p_adj_*<*0.001; Δ=0.085, 95% CI [0.082, 0.087]). Note: panels e and f use pictograms of a mouse head and neuron model to distinguish experimental and simulation data. This convention is maintained throughout the manuscript.

The *confidence c* in the reward prediction is defined as a monotonically decreasing function of the prediction uncertainty 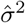, see equation (7). The confidence is a dynamic estimate of the squared fraction of variance explained (FVE^2^). The higher the FVE, i.e. the more variance in the observed outcomes can be predicted by the model, the higher is the confidence in that model and its outcome predictions. After learning, if a rule reversal takes place (Fig. 2a), highly unexpected outcomes will be observed (Fig. 2b). This leads to an increase in the uncertainty 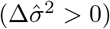, and hence a decrease in the confidence about the prediction (Δ*c <* 0, Fig. 2c). The *confidence loss* (CL) provides an internal signal, telling the animal that a change of context has occurred. The CL is elicited when unexpected outcomes are observed (see Confidence loss from unexpected outcomes). Yet, to affect behavior, the knowledge that the context has changed should take effect earlier within trial time, when sensory signals are processed and actions are decided. Neurons in the lOFC have been shown to respond to unexpected outcomes following a rule reversal [19, 29, 30]. We therefore explored whether lOFC also produced temporally advanced anticipatory signals informing that a rule reversal has occurred.

Analogously to the reward-predictor used to model top-down afferents onto sensory neurons (Fig. 1g), we modeled the lOFC activity to become predictive for the CL triggered by unexpected outcomes. In consequence, the anticipatory signal in the stimulus window predicts the CL in the outcome window. Since this anticipatory signal is reinforced by steady CL from successive unexpected outcomes, we refer to the temporal predictor of the CL as *context-prediction error* (CPE, see Context-prediction error in lOFC anticipates the confidence loss). Because the CPE is predictive for the CL that is itself based on the second-order statistics 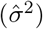 of the reward-prediction error, the CPE can be viewed as a second-order reward-prediction error. Accordingly, within few trials following a rule reversal, the CPE produced a robust internal signal of a contextual change—in this case of the fact that the rule was switched (Fig. 2c). To test this model, we analyzed calcium recordings of L2/3 lOFC neurons [19] from mice undergoing reversal learning of a go/no-go texture discrimination task (Fig. 2d). We found that the lOFC response during the stimulus window increases in the trials closely following reversal (Fig. 2e), which is consistent with a neural population encoding the CPE (Fig. 2f). Next, we investigated how the lOFC could influence learning and behavior by modeling the lOFC population with the CPE.

### Lateral OFC cancels top-down amplification in S1 after rule reversal

In scenarios where the context can change, the amplification of sensory features conditioned during a given context can be ill-suited in another context. Since lOFC neurons can signal changes in context prospectively at stimulus time (Fig. 2e), it raises the question of whether the lOFC could help to reshape top-down amplification at the apical dendrites. SST interneurons are prominent inhibitors of apical dendrites [31, 32], and OFC projections to auditory cortex have been found to be crucial for flexible auditory categorization [33]. Experimental findings have also shown that post-learning reactivation of SST could restore naïve-like activity patterns [34] and that lOFC can inhibit apical dendrites of pyramidal neurons via SST interneurons in V1 [21] and A1 [22]. We therefore presume that lOFC projections to SST interneurons in S1 counteract context-dependent top-down excitation (see Context-prediction error inhibits apical dendrites in S1 via SST interneurons, Fig. 3a and Extended Data Fig. S7). Doing so, strong lOFC responses during the stimulus windows following a rule reversal will inhibit previously conditioned sensory amplification patterns and allow the new reward-associated stimulus to be amplified.

**Fig. 3.**
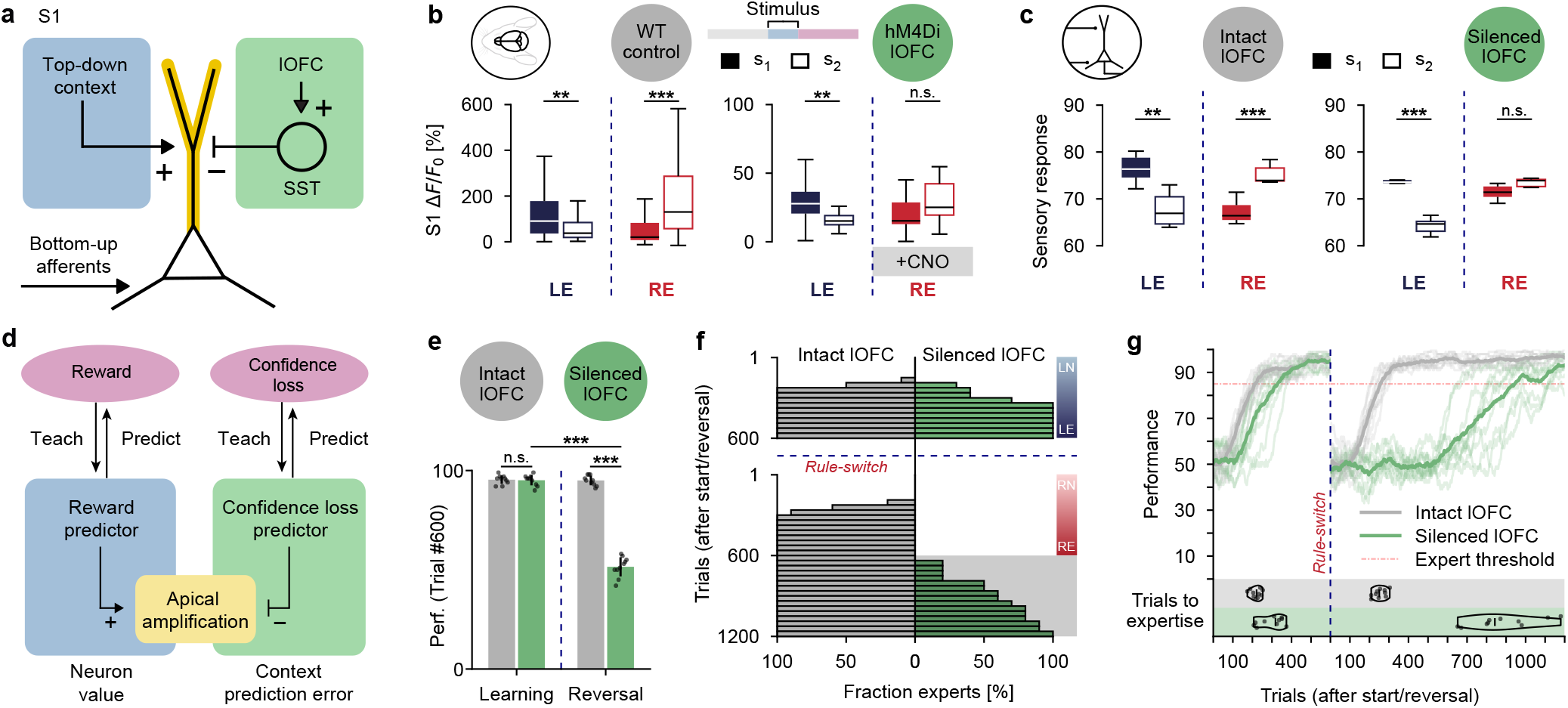
Lateral OFC inhibits the top-down amplifications in S1 upon reversal. **a**, Apical dendrites of the S1-pyramidal neurons (yellow) are excited by top-down contextual afferents (blue) and inhibited by the lOFC (encoding CPE) via SST interneurons (green). **b**, Box plots of Ca^2+^ transient amplitudes (ΔF*/*F_0_) of L2/3 S1 neurons during stimulus presentation (s_1_ or s_2_) for correct trials from wild-type (WT) control and lOFC silenced animals (50, 50 neurons in the LE, RE phase respectively from 4 WT mice and 42, 32 neurons from 3 lOFC silenced mice) during expert phases (LE, RE). Chemogenetic silencing of lOFC during the reversal phase was achieved by virally induced expression of hM4di in excitatory pyramidal neurons and daily systemic CNO injection prior to each session. Statistical differences were tested by two-sided Welch’s t-tests with Bonferroni correction, WT LE (t(67)=3.6, p_adj_=0.0011; Δ=71, 95% CI [32, 111]), WT RE (t(66)=-3.7, p_adj_*<*0.001; Δ=-160, 95% CI [-245, -75]), lOFC silenced LE (t(49)=3.0, p_adj_=0.0084; Δ=19.6, 95% CI [6.5, 32.7]), lOFC silenced RE (t(51)=-2.2, p_adj_=0.068; Δ=-17.3, 95% CI [-33.3, -1.3]). **c**, Box plots of the summed firing rates of the simulated sensory neurons, ∑_*i*_ *z*_*i*_, in response to stimulus 1 (s_1_) and stimulus 2 (s_2_) for intact lOFC LE (t(17)=3.2, p_adj_=0.0096; Δ=8.4, 95% CI [2.9, 13.9]), intact lOFC RE (t(18)=-5.0, p_adj_*<*0.001; Δ=-8.0, 95% CI [-11.3, -4.6]), silenced lOFC LE (t(13)=8.0, p_adj_*<*0.001; Δ=8.8, 95% CI [6.4, 11.2]), silenced lOFC RE (t(11)=-0.080, p_adj_=1.0; Δ=-0.13, 95% CI [-3.8, 3.5]). **d**, Functional schematic of panel a, showing the mirror-symmetry between the prediction of reward and the prediction of confidence loss. The contribution of a sensory neuron to the prediction of the future reward determines its neuron-specific value (blue). The context-prediction error (green) is a signal anticipating a confidence loss, which hence counteracts the reward-guided sensory amplification ahead of the outcome. **e**, Simulated performance (after 600 trials, RE) with and without lOFC for the original and for the reversed rule; intact vs. silenced lOFC during Learning (t(17)=0.27, p_adj_=1.0; Δ=0.3, 95% CI [-2.0, 2.6]). intact vs. silenced lOFC during Reversal (t(13)=24, p_adj_*<*0.001; Δ=43.5, 95% CI [39.6, 47.3]), Learning vs. Reversal for silenced lOFC (t(14)=24, p_adj_*<*0.001; Δ=43.6, 95% CI [39.7, 47.5]). **f**, Cumulative distributions of simulated agents that reached expert performance (85% correct) along the progression of trials; either with (left) or without (right) lOFC-mediated apical inhibition via SST. **g**, Performance traces of the agents with and without lOFC during reversal learning. Bottom: violin plots show the distributions of the number of trials necessary to become expert for the original or the reversed rule.

Our Ca^2+^ imaging data of L2/3 excitatory S1 neurons shows that the enhanced response to the rewarded stimulus s_1_, acquired during learning of the first rule, moves to the newly rewarded stimulus s_2_ after learning the reversed rule (Fig. 3b). When chemogenetically silencing lOFC neurons after the rule reversal (via CNO-activated hM4Di, see Chemogenetic silencing), the response strength between the two stimuli could no longer be distinguished significantly. The evolution of the S1 response and its dependence on lOFC could be reproduced in the model by including a lOFC feedback to SST interneurons of the sensory cortex (Fig. 3c; Supplementary Methods 3). With this inhibitory feedback, the selective top-down amplification of reward-predictive sensory stimuli can be reverted by lOFC, provided that a context change leads to a strong context-prediction error (Fig. 3d).

Besides the sensory responses, we also compared the performances achieved by the model with and without lOFC after reversal. We found that the model without lOFC failed to adapt to the reversed rule within 600 trials (Fig. 3e). However, if the context-prediction error from lOFC drives apical inhibition in S1, the incorrect amplification of sensory neurons from bygone contexts was prevented. In consequence, by unbiasing the sensory representation, lOFC allows the reversed rule to be learned almost as fast as the original rule (Fig. 3f). Interestingly, animals with silenced lOFC can eventually adapt to the reversed rule after significantly more trials Extended Data Fig. S9, which the model also reproduced (Fig. 3g).

### Context-prediction errors support learning for multiple reversals

Besides unbiasing the sensory representation, inhibiting the top-down sensory amplification upon rule reversal also directly affects action selection (Fig. 4a). The conditioned behavior is inhibited at the source, by reducing the response to the stimuli responsible for triggering the behavior in the first place. At the same time, reducing sensory neuron responses also decreases the plasticity of their efferent synapses (see Supplementary Equations 1). As a consequence, the synapses encoding the policy for the original rule are protected from degradation, which becomes useful in case of a second reversal back to the original rule (Fig. 4b). In other words, behavior can be adapted to the reversed rule, while still preserving the synaptic weights encoding the sensorimotor mapping for the original rule (Fig. 4c and Extended Data Fig. S10). Upon a second rule reversal, we observe that the mice adjusted their behavior faster than during the first reversal (Fig. 4d). Within the model, this can be attributed to the protection of the sensorimotor mapping acquired in the first learning stage. Indeed, behavioral adaptation requires both an appropriate sensorimotor mapping and sensory top-down modulation [7] (Fig. 4a). As only the latter needs to be adjusted after the second reversal (Fig. 4e), the overall behavioral adaptation can occur more rapidly.

**Fig. 4.**
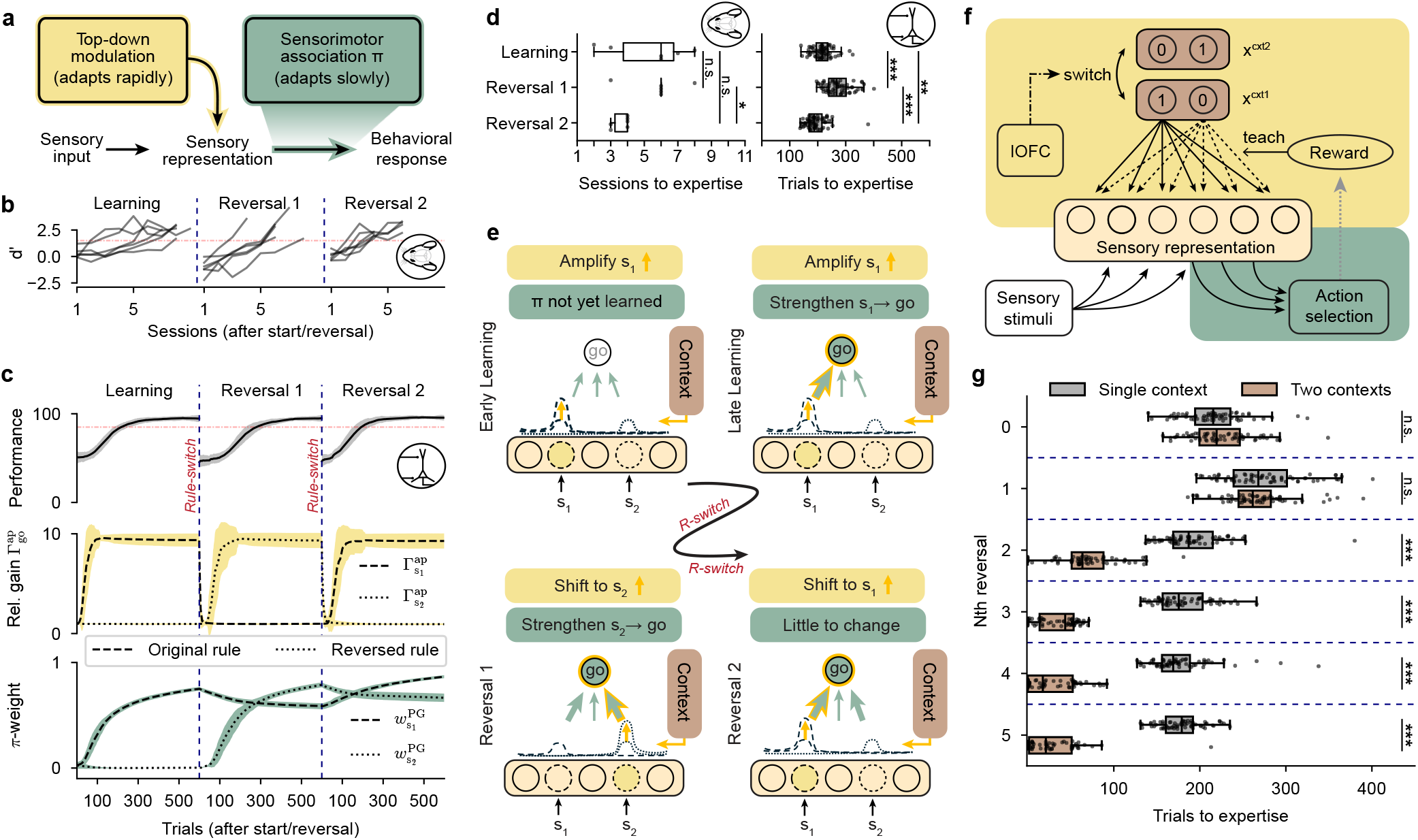
The context-prediction error speeds up adaptation following multiple reversals. **a**, The two main components involved in behavioral learning: rapidly adapting top-down modulation of the sensory representation depending on the context (yellow), and slower-adapting sensorimotor association *π* (green). **b**, Performance traces of 6 mice during two consecutive rule reversals, where the second one reverts back to the original rule. **c**, Top: Average performance of the agents for two sequential reversals. Middle: Evolution of the relative gain, see equation (12), of the go-stimulus-selective neurons, which corresponds to the s_1_-selective neuron when the original rule applies (dashed) and to the s_2_-selective neuron for the reversed rule (dotted). After each reversal, the top-down inhibition rapidly erases the relative gain. Bottom: The *π*-weight represents the synaptic strength of the weight in the policy network connecting from the go-stimulus-selective neuron to the go-action neuron. After potentiating the synaptic weight from the s_1_ neuron to the go action during Learning, it remains high throughout Reversal 1, while the synaptic weight from the s_2_ neuron to the go action is strengthened. **d**, Number of sessions needed for each of the 4 mice to reach expertise for Learning, Reversal 1, and Reversal 2 (left); Learning vs. Reversal 1 (Log-rank: *χ*^2^(df=1)=0.045, p_adj_=1.00000; Cox PH: HR=0.98, 95% CI [0.31, 3.04]), Learning vs. Reversal 2 (Log-rank: *χ*^2^(df=1)=2.45, p_adj_=0.35; Cox PH: HR=0.25, 95% CI [0.05, 1.28]), Reversal 1 vs. Reversal 2 (Log-rank: *χ*^2^(df=1)=6.02, p_adj_=0.042; Cox PH: HR=0.10, 95% CI [0.012, 0.79]); and the same in trials for the model (right); Learning vs. Reversal 1 (Log-rank: *χ*^2^(df=1)=51, p_adj_*<*0.001; Cox PH: HR=3.8, 95% CI [2.6, 5.5]), Learning vs. Reversal 2 (Log-rank: *χ*^2^(df=1)=10.4, p_adj_=0.0038; Cox PH: HR=0.57, 95% CI [0.40, 0.81]), Reversal 1 vs. Reversal 2 (Log-rank: *χ*^2^(df=1)=75, p_adj_*<*0.001; Cox PH: HR=0.20, 95% CI [0.13, 0.29]). **e**, Qualitative illustration of the steps that led to faster adaptation after the second reversal. Early learning: amplification of the s_1_-selective sensory neuron. Learning: increasing the synaptic weight from the s_1_-selective neuron to the go-action neuron. Reversal 1: top-down sensory amplification is shifted from the s_1_ to the s_2_-selective sensory neuron, and the weight from the s_2_-neuron to the go-action neuron is strengthened. Reversal 2: synaptic weights from the s_1_-selective neuron to the go-action neuron remain strengthened; only the sensory modulation needs substantial adjustment. **f**, Model in which two different contexts can be represented and where the context representation *x*^ap^ can be switched by lOFC activity (context-prediction error). Allowing the sensory modulation to be adapted by changing *x*^ap^ removes the need to re-learn the top-down modulation weights in the case when a previously learned context reappears. **g**, Number of trials necessary to reach expert performance with a single context representation (gray) and with two context representations (brown) for five sequential rule reversals. All statistical differences were tested using a two-sided log-rank test with Bonferroni correction and effect size was estimated with a Cox proportional hazards model; Learning (Log-rank: *χ*^2^(df=1)=1.92, p_adj_=0.99; Cox PH: HR=1.28, 95% CI [0.90, 1.82]), Reversal 1 (Log-rank: *χ*^2^(df=1)=2.15, p_adj_=0.86; Cox PH: HR=0.77, 95% CI [0.54, 1.10]), Reversal 2 (Log-rank: *χ*^2^(df=1)=145, p_adj_*<*0.001; Cox PH: HR=0.050, 95% CI [0.029, 0.087]), Reversal 3 (Log-rank: *χ*^2^(df=1)=155, p_adj_*<*0.001; Cox PH: HR=0.0061, 95% CI [0.0014, 0.026]), Reversal 4 (Log-rank: *χ*^2^(df=1)=158, p_adj_*<*0.001; Cox PH: HR=0.0061, 95% CI [0.0014, 0.026]), Reversal 5 (Log-rank: *χ*^2^(df=1)=122, p_adj_*<*0.001; Cox PH: HR=0.10, 95% CI [0.067, 0.16]).

By preserving the sensorimotor mappings of prior rules, the appropriate behavior for these rules can be restored merely by recovering the top-down sensory amplification pattern. Until now we considered top-down excitation of sensory neurons to be driven by a single, fixed context representation vector. Given the single context representation, the top-down projections had to re-learn the contextual value of each neuron upon each rule switch (Fig. 4c). To that end, the synaptic weights exciting formerly amplified neurons had to be weakened; and different synapses had to be strengthened. An alternative strategy is to modify the sensory amplification by switching the context representation while maintaining the synaptic weights that transmit the top-down excitation [14] (Fig. 4f). It has been shown that sensory amplification is context-dependent [6, 23, 35, 36] and that animals can develop dedicated task-representing neurons [37]. Furthermore, activity in the human lOFC has been suggested to represent a shift in context representation [29]. We, therefore, explored whether this principle would yield computational benefits (see lOFC triggers switch of context representation). For that, we simulated agents with two possible context representations, each consisting of a two-dimensional one-hot vector and projecting to the apical dendrites of the sensory neurons. The lOFC activity encoding the context-prediction error was then used to trigger a switch between context representations (Fig. 4f). The capacity to represent different contexts and to switch between these representations significantly improved behavioral adaptation following multiple rule reversals (Fig. 4g). Indeed, when a rule had already been experienced previously, not only were the stimulus-to-action projections already learned (as in the single-context model), but also the top-down excitation weights onto the sensory neurons (Extended Data Fig. S11a). While we implemented the two-context model, the lOFC to sensory cortex SST interneuron feedback became less important, since the context-representation switch prevented the inadequate sensory amplification from the prior rule. Nonetheless, SST interneuron activation could provide a necessary inhibition in case the new context-representation is not yet established or if top-down signals from other sources (e.g. motor regions) must be suppressed following reversal. Although lOFC triggering a switch in contextual representation offers an appealing account of behavioral adaptation across multiple reversals, it is still too early to draw a definitive conclusion. Experiments to test predictions derived from each mechanism are discussed in the Discussion.

### VIP-mediated disinhibition reinforces anticipatory sensory signals

We showed how top-down signals such as the context-prediction error could improve sensory processing to accommodate different behavioral contexts. While we provided a mathematical framework, the neural mechanism by which the lOFC learns to generate the context-prediction error signal during the stimulus window is still unclear. In general, through the accumulation of experience across trials, anticipatory responses build up ahead in time from the reinforcers they learn to predict. While a variety of such predictive signals can be observed in the cortex [38, 39], little is known about the underlying microcircuits and neural mechanisms that generate them. Accumulating evidence shows that VIP-mediated disinhibition [40–42] may play a critical role in gating associative learning in the olfactory [43], auditory [44], visual [45, 46] and the somatosensory cortex [47], and could potentially be part of a cortex-wide mechanism [48]. In different cortical regions, VIP interneurons are recruited by behavioral reinforcers such as reward and punishment [49]. Hence, VIP interneurons could play a role in signaling to pyramidal neurons that a salient outcome has occurred and gate the emergence of a predictive response. Applying these ideas to the lOFC model, we suggest that a population of lOFC VIP interneurons could convey the confidence loss (CL) to CPE encoding lOFC neurons via disinhibition (Fig. 3d). This hypothesis predicts that lOFC signaling CPE depends on lOFC VIP interneurons. In consequence, reversal learning would be degraded by silencing lOFC VIP interneurons, which can be simulated in the model by setting CL = 0 (Fig. 5a).

**Fig. 5.**
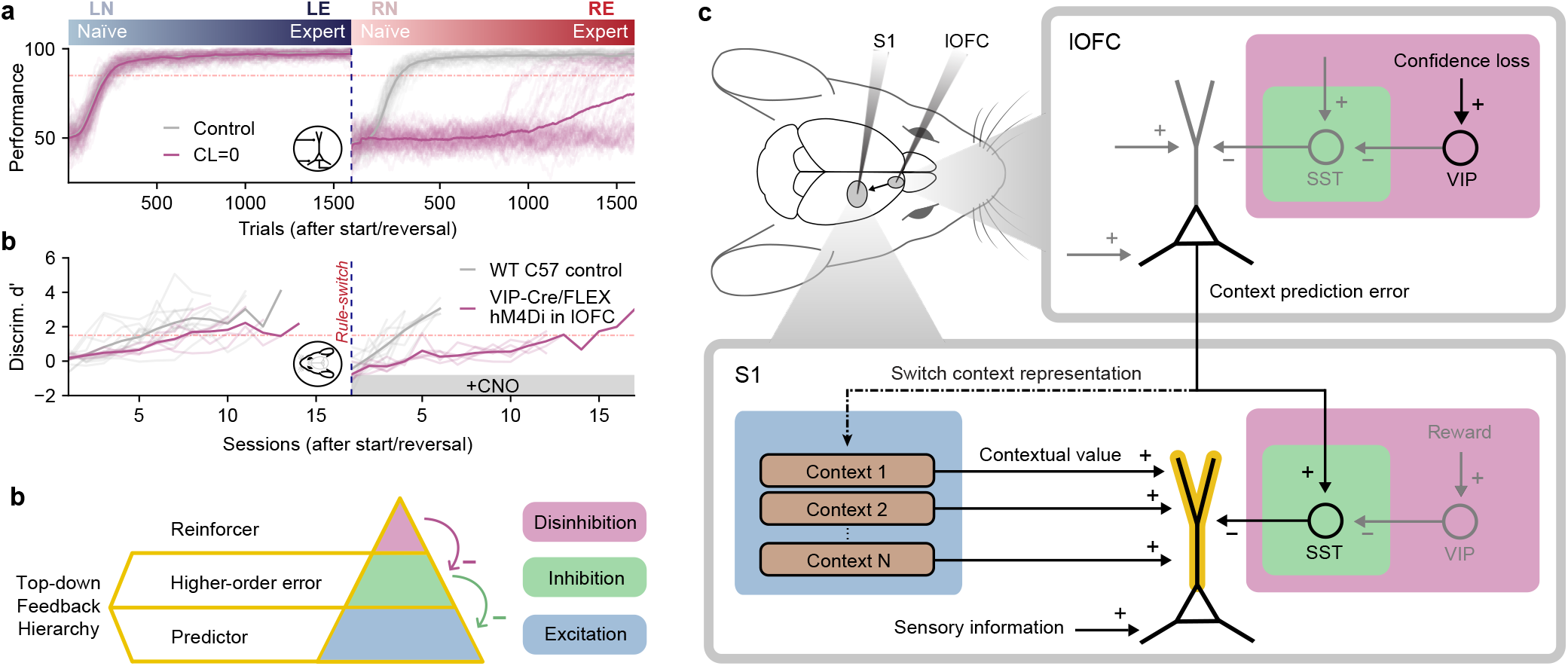
VIP interneurons as local reinforcers for learning a hierarchy of anticipatory signals. **a**, Top: Simulated performances during reversal learning with the confidence loss (CL) fixed to zero (gray) compared to the undisturbed control case (purple) Bottom: Performances of mice with (purple) and without (gray) chemogenetic silencing of their lOFC VIP neurons (10 and 5 mice, respectively). Statistical analysis comparing the silenced VIP and the control condition are reported in Extended Data Fig. S12. **b**, Top-down feedback hierarchy onto pyramidal neurons, where inhibition overrules excitation, and disinhibition overrules inhibition. At the lowest level (blue), the top-down excitation targeting the apical dendrites of pyramidal neurons represents a context-dependent anticipatory predictor (in S1 of reward, in lOFC of a confidence loss). At the middle level (green), the SST-mediated top-down inhibition counteracts the lower-level excitation by a higher-order error (in S1, the context-prediction error counteracting the neuron-specific apical value). At the top level (purple), the top-down disinhibition represents a reinforcer conveyed by VIP interneurons (in S1 an action-induced reward signal, in lOFC the confidence loss). **c**, Microcircuits implementing the top-down feedback hierarchy from b within S1 and lOFC, with apical excitation (blue) conveying contextual value in S1 that is inhibited by the context-prediction error signal (green), and a new anticipatory context signal (e.g. Context 2 in S1) being reinforced through disinhibition via VIP interneurons (purple).

To test this hypothesis *in vivo*, we silenced VIP interneurons by virally inducing hM4Di expression using a Cre-dependent viral construct in VIP-Cre mice and administering intraperitoneal CNO during reversal learning (RN through RE phase). Silencing lOFC VIP interneurons, significantly delayed reversal learning (Fig. 5a, Extended Data Fig. S12). The model suggests that this deficit can be attributed to an impairment in the reorganization of the sensory representation upon reversal. In the control condition, the model uses lOFC activity to drive sensory SST neurons, as experimentally observed in V1 [21] and A1 [22]. If VIP interneurons in lOFC are silenced, the confidence loss triggered during the outcome window is prevented from reinforcing the anticipatory context-prediction error during the stimulus window (Fig. 2c). As a consequence, the previously learned apical values in S1 pyramidal neurons are not suppressed by top-down inhibition upon the rule-reversal (Fig. 5b,c). In S1, the previously reward-predictive sensory neurons remain enhanced and learning is disturbed, because the action selection network has to compensate for the incorrectly modulated sensory representation (see also Fig. 4e).

## Discussion

We performed calcium imaging experiments in S1 and lOFC of mice during a sensory learning task with subsequent reversal of the association rule. While learning the initial rule, the sensory response in S1 to the reward-predictive stimulus was enhanced. After reversing the rule, the stimulus enhancement to the previously reward-predictive stimulus was decreased, while the response to the reward-predictive stimulus in the second rule was enhanced. In lOFC, calcium responses signaled the occurrence of a rule-switch and lOFC silencing impaired the reorganization of the sensory representation in S1. A theory is presented that explains the role of lOFC in reversal learning by reshaping the top-down modulation of S1 responses after a rule switch. The theory is supported by a computational model with components constrained by existing experimental findings ranging from sensory response adaptation [5–7, 9– 11, 19, 23, 26–28, 35, 36], somatic excitability changes from apical dendrite activity [13, 24], apical dendrites responses [15, 16, 25], lOFC neuron responses [19, 29, 30], the role of lOFC in reversal learning [19, 20, 53], the inhibition of apical dendrites by SST interneurons [31, 32], task-encoding in prefrontal cortex neurons [37], the disinhibition effect of VIP interneurons [40–42], to the role of VIP interneurons in associative learning [43–48].

To explain the key features of the experimental observations, the computational model makes use of the notions of confidence, confidence loss, and context-prediction error. During learning, the confidence in the predicted outcome increases. An unexpected rule-switch results in a confidence loss at the moment when the previously correct outcome prediction turns out to be wrong. Such a confidence loss is signaled by lOFC after the rule switch, when the expected reward during the outcome window is missed. Crucially, in trials following the rule-switch, lOFC advanced its firing in trial time and learned to signal a context-prediction error during the stimulus window, being predictive for the confidence loss in the outcome window. In S1, the sensory response to the reward-predicting stimulus is enhanced during initial learning. The stimulus enhancement is neuron-specific and proportional to the contribution of the neuron to the reward prediction. With the signaling of the context-prediction error by the lOFC after the rule-switch, the response enhancement in S1 is undone via apical inhibition through local SST interneurons. During the subsequent re-learning, the sensory response to the newly reward-predictive stimulus becomes enhanced through a reorganization of the top-down input to S1. Blocking experiments suggest that VIP interneurons in lOFC signal the confidence loss to the lOFC pyramidal neurons, providing the reinforcement signal to develop the anticipatory context-prediction error. This context-prediction error facilitates the reorganization of the top-down induced gain modulation in S1.

The behavior-dependent reorganization of sensory representation during learning is well recognized [50–53], and top-down feedback has been established as a fundamental component of perceptual learning [9, 11, 12, 14, 16, 17, 27]. Our work revealed how early regions of sensory processing can be modulated by lOFC to reshape sensory computation depending on task context. The importance of lOFC in reversal learning and contextual response-outcome associations has been well documented in mice [19, 53], rats [54], non-human primates [55], and humans [20, 56]. We recently reported that similar neural circuits involving lOFC could be operational using analogous cognitive strategies in distinct species [57, 58]. While many different roles have been proposed for the lOFC to support cognitive flexibility [59, 60], its importance has been especially highlighted for state representation learning [61–63], and for representing confidence [64, 65] and context switching [29]. The proposed computational model provides a unifying account in which lOFC signals changes in task context and supports the reorganization of the sensory representation.

The hierarchy of reward- and context-prediction errors can be viewed as a mechanism to reduce different sources of uncertainties. Selective amplification prevents irrelevant information from disturbing the learning process, which reduces the expected uncertainty [66] as long as the context remains the same. While uncertainty estimates have been suggested for global learning-rate modulation [67], regulating selective top-down amplification provides a mechanism for a neuron-specific learning-rate modulation (see Supplementary Equations 1). A rule reversal elicits uncertainty about the current context, which can be termed unexpected uncertainty (see Supplementary Equations 3), and should reduce reliance on top-down contextual information [66]. SST interneurons have been shown to drive a shift away from top-down influences to favor bottom-up information streams [34]. OFC projections to SST interneurons in S1 provide a direct mechanism to inhibit selective top-down amplification and unbias the sensory representation in case of contextual uncertainty. This offers a mechanistic basis for cortical learning under uncertainty, which is distinct from learning through precision-weighted prediction errors [68]. The unexpected uncertainty signaled by the context-prediction error in the lOFC directly modulates sensory neuron responses and, consequently, action selection probabilities. By contrast, precision-weighted plasticity relies on estimates of expected uncertainty to modulate synaptic weight updates, thereby affecting future neuronal responses and behavior only indirectly.

Disinhibition by VIP interneurons represents a local reinforcement signal that varies depending on the cortical region [47, 49, 69]. In the basolateral amygdala, VIP interneurons respond strongly to punishments and are necessary to develop enhanced responses to conditioned stimuli during fear learning [70]. VIP interneurons are also directly affected by neuromodulators. In cortex, the only ionotropic serotonin receptors (5-HT3R) expressed are found in interneurons that do not express PV or SST [71] and co-express VIP and CCK [72]. Converging evidence indicates that serotonergic signaling in OFC is critical for reversal learning [73–75]. Serotonin has been associated with sensory uncertainty [76], unsigned reward-prediction error [77], surprise [78], and expected and unexpected uncertainty [67]. Our hypothesis that lOFC VIP interneurons represents confidence loss (Fig. 5c) is compatible with these reports, offers a potential explanation for the role of lOFC serotonin in reversal learning, and provides a testable framework to guide future experiments.

The proposed hierarchical model of context-dependent sensory processing is strongly supported by biological evidence, but also makes assumptions that call for further experimental confirmation. (*i*) There is strong support for a reward-associated top-down modulation of the sensory representation (Fig. 1g); however, the precise biological mechanisms coordinating this modulation remain unclear. It requires understanding how the brain solves the credit-assignment problem, which is still an open area of investigation. (*ii*) While lOFC has been shown to target SST interneurons in V1 [21] and A1 [22], explaining the S1 recordings with the lOFC-to-SST interneurons pathway presumes that these projections also exist in S1 (Fig. 3a). (*iii*) The second proposed mechanism for the lOFC to modulate the sensory representation suggests that the context-prediction error induces a change in context representation (Fig. 4f). Task-encoding prefrontal cortex neurons are plausible suspects for this mechanism, but demonstrating how the lOFC might trigger a shift in the context representation still requires further investigation. (*iv*) Predictive reinforcement through disinhibition provides a potential explanation for the emergence of reward-associated amplification in sensory neurons and the context-prediction error in the lOFC, as well as the dependence of reversal learning on lOFC VIP interneurons, but this mechanism remains speculative.

Besides highlighting gaps in experimental knowledge, the theory also makes testable predictions that can instruct future experiments. (a) It predicts context-dependent afferents that modulate the responses of sensory pyramidal neurons according to task-relevance. (b) The two proposed mechanisms for the lOFC to reshape the sensory representation can be tested behaviorally. If reversal learning only depends on the lOFC-to-SST feedback (Fig. 3a), a strong context-prediction error response in lOFC will be required following each reversal. If lOFC directly changes the context representation (Fig. 4f), the lOFC response upon a rule reversal would decrease for subsequent reversals (Extended Data Fig. S11b). (c) The lOFC-to-SST feedback mechanism also predicts that the context-prediction error should be reflected in the activity of sensory SST interneurons and that inhibiting direct lOFC projections to sensory cortex would impair reversal learning. (d) Finally, the hypothesis proposing that VIP interneurons drive the context-prediction error predicts that the confidence loss should be encoded in VIP interneuron activity in the lOFC. These predictions should be tested in future studies and offer concrete steps to assess the proposed circuit mechanisms.

The richness with which the brain modulates sensory representations through first- and second-order reward-prediction errors could also have applications in machine learning. Dendritic nonlinearities were suggested to increase the expressive power of neural networks [79] and context-dependent dendritic gain modulation was proposed as a biologically plausible mechanism for multi-task learning and as a means to reduce catastrophic forgetting in continual learning [14, 37, 80–82]. However, these models lacked an explanation for how the brain can recognize a change of context and signal the context change during sensory perception, before a behavioral decision is made. Prospective errors were suggested to project to apical dendrites of pyramidal neurons and prospectively correct the activity of deep neurons in the cortical hierarchy [83–86]. The presented results on context-prediction errors are in line with such normative theories and offer a framework for understanding the biological substrate of context-dependent computation that may inspire future progress in both neuroscience and artificial intelligence [3, 87].

## Methods

### Gain modulation in sensory representations

The sensory representation ***z*** consists of *N* pyramidal neurons *i* ∈ {1, …, *N*} with two compartments. The basal compartment received bottom-up sensory inputs with fixed selectivity, producing activations ***x***^bas^, and the apical compartment dictated the multiplicative gain ***g***^ap^ of the firing rates,

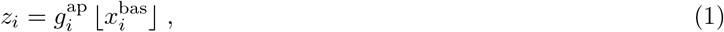

where ⌊·⌋ stands for the rectifier function ⌊*y*⌋ = max(0, *y*). An apical compartment receives *N*_cxt_ = 2 context-dependent excitatory inputs 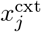, weighted by the synaptic strength 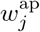. It also receives global inhibition *r*^SST^ from a population of SST interneurons, see equation (10). These top-down apical afferents determine the context-specific apical gain 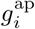 within the range [1, *g*_max_],

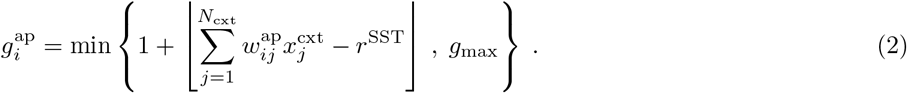

The *salience* of a sensory stimulus is proportional to the gain 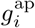 of its sensory neurons. The gain is increased by apical excitation 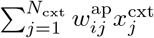 that originates in the *context representation* ***x***^cxt^. The apical excitation can be canceled by the SST interneuron activity *r*^SST^, or altered by changing ***x***^cxt^ (as in Fig. 4f).

### Apical amplification based on neuron-specific value

The apical weights ***w***^*ap*^ are learned to amplify sensory neurons that are predictive for reward. Each neuron is assigned a neuron-specific value *v*_*i*_, encoding how much this neuron contributes to the estimate of the (global) value *V*(*z*) = E[*R* | ***z***], i.e. to the expected future reward conditioned on the current state representation. The value estimator 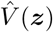 is calculated based on the sensory representation ***z*** as

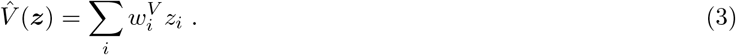

The weights 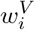 are adapted to reduce the reward-prediction error 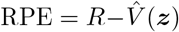, constrained by a L1 regularization,

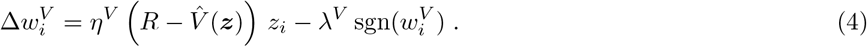

This weight update reduces the loss *L*^*V*^ = E[RPE^2^] + *λ*^*V*^ ∥***w***^*V*^ ∥_1_, where ∥.∥_1_ refers to the L1-norm. The neuron-specific value *v*_*i*_ encoding how much a neuron *i* predicts an upcoming reward is defined by its non-negative contribution to the value estimation 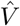, i.e. 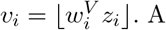 key ingredient of the model is that the sensory representation is sharpened by the top-down input, such that the task-relevant sensory neurons are amplified compared to task-irrelevant neurons. The criterion for task-relevance is set by the comparison between the value *v*_*i*_ of a neuron and the activity *r*^SST^ of the SST interneurons. If *v*_*i*_ *> r*^SST^, the apical excitation is enhanced. If *v*_*i*_ *< r*^SST^, the neuron *i* is not sufficiently relevant and its apical excitation is reduced. This yields the learning rule for the excitatory apical synapses

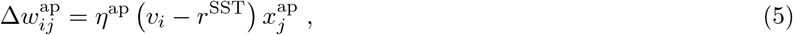

where *η*^ap^ represents the learning rate and 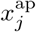 is the presynaptic activity, see equation (2). Asymptotically, this plasticity rule leads to a binary distribution of the apical weights. Active synapses projecting to a highly task-relevant sensory neuron become potentiated and converge towards 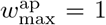. Synapses projecting to task-irrelevant neurons become depressed and converge towards 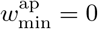 (see Supplementary Equations 2). Taken together, the RPE informs the value estimate and hence the gain amplification of individual L2/3 sensory neurons.

### Estimating the confidence of the reward prediction

In order for the brain to notice a change of context, it first needs a notion of understanding the current context at hand. The action-value estimate 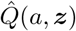 approximating E[*R*|*a*, ***z***]—while also supporting action selection (see Supplementary Methods 2)—encodes the current understanding of the agent about the task. The action-value estimate 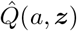 is computed from an outcome prediction network that implicitly represents the context, i.e. the task rule, through neurons becoming selective for stimulus-action-outcome triples (see Supplementary Methods 1).

The variance unexplained (VU) by the model can be measured by the expected squared reward-prediction error 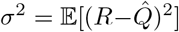, see Supplementary Equations 4. The variance unexplained, *σ*^2^, jointly measures the bias in the reward prediction, its distraction by task-irrelevant stimuli, and the intrinsic variability of *R* (see Supplementary Equations 3). We assume that the brain approximates *σ*^2^ with an internal estimate that we denote as *prediction uncertainty* 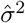[88]. The prediction uncertainty 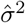 is updated based on the difference between the current squared reward-prediction error at the current time step and the uncertainty estimate in the previous trial,

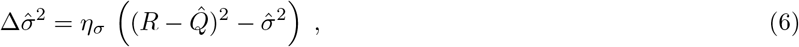

with a small learning rate *η*_*σ*_. If 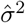 is small, the animal can be confident in its model of the task and the reward prediction. We therefore defined the *prediction confidence c* as estimate of the squared *fraction of explained variance* (FVE), or also squared coefficient of determination (see Supplementary Equations 4),

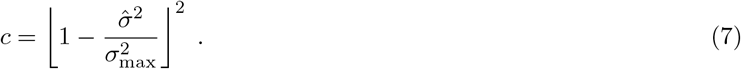

Here, 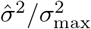 represents the fraction of variance unexplained (FVU). The maximal variance 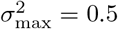 was computed for a uniform random policy (see Supplementary Equations 5). The prediction confidence *c* is restricted to values between 0 and 1. For more details see Supplementary Equations 4.

### Confidence loss from unexpected outcomes

The prediction uncertainty 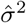 can also be understood as a form of expected uncertainty given the internal model of the task [66, 89]. The prediction confidence, correspondingly, can be seen as model confidence. When an outcome occurs that lies further away from the prediction than would be expected, it can elicit unexpected uncertainty [67]. Translated to our framework, if 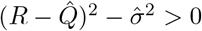, the brain signals a higher-order error compared to the first-order prediction error 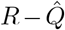. As the rat OFC has been shown to be involved in confidence computation [65], we suggest that the *confidence loss* (CL) could also be signaled to lOFC. In the model, CL is defined as decrease in model confidence induced by the change 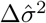 given in equation (6), normalized by the learning rate *η*_*σ*_. The confidence loss can be expressed as a function of 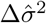,

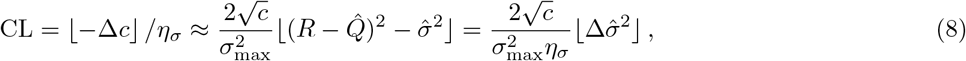

see Supplementary Equations 6. Hence, the CL can be viewed as a confidence-weighted 2nd-order error of the reward prediction. A strong confidence loss is contingent on the animal being confident in its model of the environment (*c >* 0).

### Context-prediction error in lOFC anticipates the confidence loss

We denote the anticipation of the CL at stimulus time the *context-prediction error* (CPE). While the confidence loss CL is elicited when an unexpected outcome is observed during the reward presentation window *t*_R_, the contextual uncertainty can be represented in the next trial already during the stimulus window *t*_S_. The CPE is learned by minimizing the squared difference between CL and CPE, 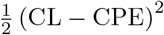. Minimizing this squared difference online by gradient descent leads to an update of the CPE by

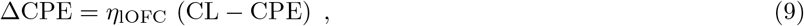

with learning rate *η*_lOFC_. In the simulations, we model the lOFC activity during the stimulus window as context-prediction error, *r*^lOFC^ = CPE.

### Context-prediction error inhibits apical dendrites in S1 via SST interneurons

The SST interneurons in S1 are driven by the context-prediction error encoded in lOFC. They modulate the sensory processing in two ways. First, they directly reduce the gain of the sensory representation neurons in S1 by inhibiting the apical dendrite activity, see equation (2). Second, they represent a task-relevance threshold that the neuron-specific value *v*_*i*_ has to reach in order to potentiate excitatory apical synapses, see equation (5) and Supplementary Equations 2. The firing rate of the SST population in S1 at the time of stimulus presentation *t*_S_ was modeled as

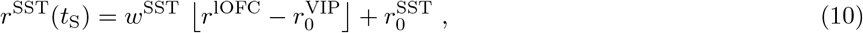

where 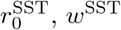 and 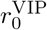 are constants.

### lOFC triggers switch of context representation

In most simulations, the gain amplification conveyed by excitatory apical synapses ***w***^ap^ originated from a single, fixed context representation ***x***^cxt^ = (1, 0). For this single-context model of the brain, the apical gain could only be adapted by changing the synaptic weights ***w***^ap^ or by tuning the inhibitory contribution from SST interneurons. Alternatively, different contexts representations ***x***^cxt^ can be considered, and the gain can be modified by switching the context. To show this, we implemented context 1 by ***x***^cxt1^ = (1, 0) and context 2 by ***x***^cxt2^ = (0, 1). The switch between both contexts was triggered when the lOFC activity reached a threshold activity *θ*_switch_ and the model confidence *c* was above a threshold *c*_switch_.

### Confidence-weighted action selection

During reversal learning, the policy depended on the sensory representation ***z*** and on the model-confidence *c*. The probability of selecting an action *π*(*a*|***z***, *c*) consisted of a combination of two separate action selection policies: one explorative policy *π*^Q^ based on action-values and one exploitative policy *π*^PG^ based on policy gradient (Extended Data Fig. S6a)

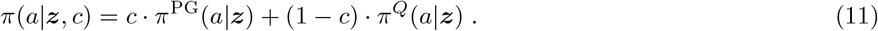

Hence, at low confidence *c*—in case of high uncertainty 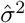—action selection was mainly driven by the explorative policy *π*^Q^. At high confidence *c*—in case of low uncertainty—the action selection was dominated by the exploitative policy *π*^PG^. For more details on the action selection (see Supplementary Methods 2, 3 and Extended Data Table 1).

**Table 1.**
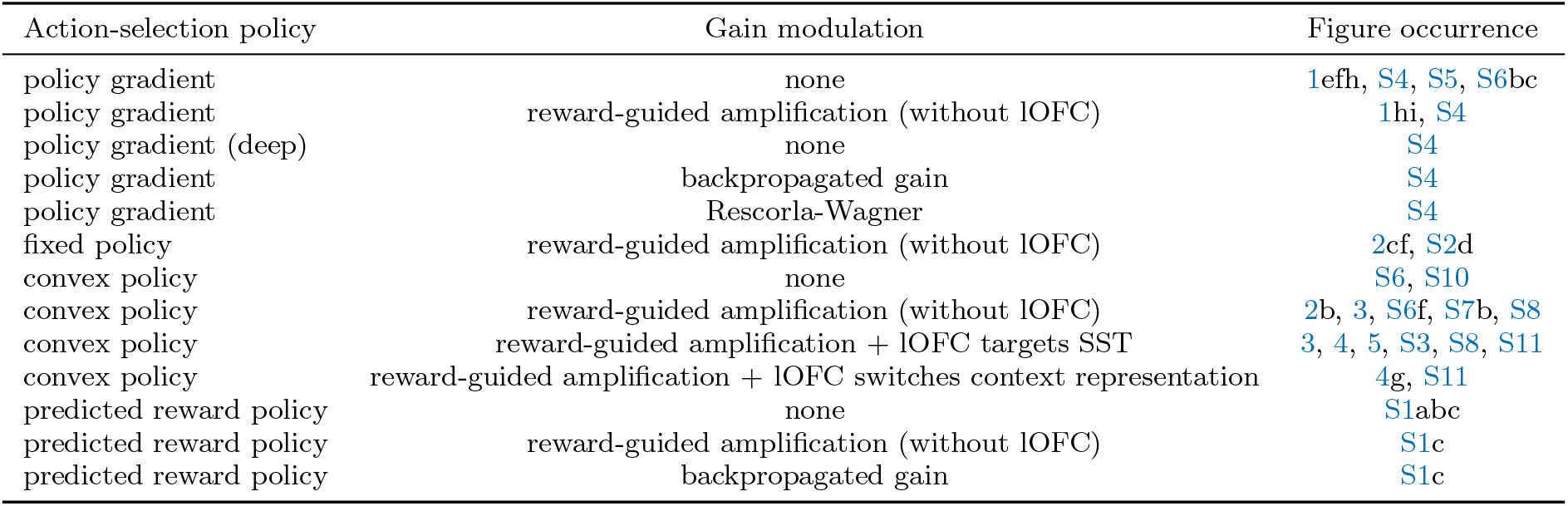
Model combinations.

### Simulated go/no-go task

Learning of the go/no-go sensory discrimination task was simulated for 600 trials (Fig. 1b). After that, the simulation was either terminated or the stimulus-reward contingency was reversed. Each trial began by randomly sampling the *N* -dimensional stimulus pattern ***x***^bas^ characterizing the basal input to the *N* sensory pyramidal neurons. In all simulations, one sensory representation neuron was exclusively selective to stimulus 1 (s_1_), which means that its basal activation 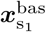 was 1 in case the stimulus 1 was presented and 0 otherwise. Analogously, one neuron was exclusively selective to stimulus 2 (s_2_). The remaining *N*_*d*_ = *N* −2 ‘distractor’ neurons received random basal inputs independently sampled from binary noise *ξ*_*i*_ ∈ {0, 1} with probability 0.5 for each trial (see Fig. 1c). In other words, 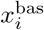 was either 1 or 0 (except for Fig. 1e, 1f, Extended Data Fig. S6d and S6e, where the s_1_- and s_2_-dependent response strengths were varied). Unless specified otherwise, we used *N*_*d*_ = 128 and *N* = 130.

The bottom-up activations ***x***^bas^ were processed to compute the sensory representation ***z***, see equation (1). Based on ***z***, the policy *π* then selected between two actions: go (*a*_GO_) or no-go (*a*_NG_). Depending on the stimulus and action combination, an outcome with reward *R* ∈ *{R*_punish_, 0, *R*_reward_} was fed back to the agent (Extended Data Fig. 1b).

### Simulation parameters

All simulations were run with 64 different pseudorandom seeds, which were all used to compute the means and other statistics displayed in the figures. When applicable, the parameters 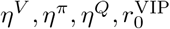and *w*^SST^ were set by a grid search algorithm (Extended Data Algorithm 2) to either maximize the performance or minimize the number of trials necessary to reach expert performance (Extended Data Table 3). All other parameters 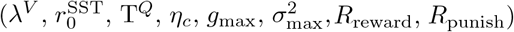 were predefined constants (Extended Data Table 2). A detailed description of the parameter settings is included in Supplementary Methods 3 and 5.

**Table 2.** Fixed parameter values.

| Parameter description | Mathematical symbol | Parameter value |
| --- | --- | --- |
| Maximal apical gain | $g_{\max}$ | 10 |
| Reward value | $R_{\text{reward}}$ | 1 |
| Punishment value | $R_{\text{punishment}}$ | -1 |
| L1 regularization factor | $\lambda^V$ | 1/N |
| Excitatory apical weight learning rate | $\eta^{\text{ap}}$ | 0.1 |
| Maximal excitatory apical weight | $w_{\max}^{\text{ap}}$ | $g_{\max} - 1 = 9$ |
| Minimal excitatory apical weight | $w_{\min}^{\text{ap}}$ | 0 |
| Uncertainty learning rate | $\eta^{\sigma}$ | 0.03 |
| IOFC learning rate | $\eta^{\text{IOFC}}$ | 0.03 |
| SST interneuron baseline activity | $r_0^{\text{SST}}$ | 0.1 |
| Q-value policy temperature | $T^Q$ | 10 |
| Minimal policy exploration | $\epsilon_{\phi}$ | 0.01 |
| Policy sigmoid bias | $b_{\phi}$ | 99 |
| Maximal uncertainty | $\sigma_{\max}^2$ | 0.5 |
| CPE threshold for context switch | $\theta_{\text{switch}}$ | 1 |
| Confidence threshold for context switch | $c_{\text{switch}}$ | 0.95 |

**Table 3.** Optimized parameter values.

| Model description | $\eta^{\pi}$ | $\eta^V$ | $\eta^Q$ | $r_0^{\text{VIP}}$ | $w^{\text{SST}}$ | Optimized for | Figure occurrence |
| --- | --- | --- | --- | --- | --- | --- | --- |
| Policy gradient without gain | M | N/A | N/A | N/A | N/A | Learning perf. | 1efh, S4, S5 |
| Policy gradient with gain (without IOFC) | M | M | N/A | N/A | N/A | Learning perf. | 1hi, S4 |
| Reward prediction without gain | N/A | N/A | M | N/A | N/A | Learning perf. | S1 |
| Reward prediction with gain | N/A | M | M | N/A | N/A | Learning perf. | S1c |
| Reward prediction with backprop. gain | N/A | N/A | M | N/A | N/A | Learning perf. | S1c |
| Policy gradient with bottom-up plasticity | M | N/A | N/A | N/A | N/A | Learning perf. | S4 |
| Policy gradient with backprop. gain | M | N/A | N/A | N/A | N/A | Learning perf. | S4 |
| Policy gradient with RW gain | M | M* | N/A | N/A | N/A | Learning perf. | S4 |
| Fixed policy with gain (without IOFC) | N/A | $10^{1.25\dagger}$ | $10^{-1.5\dagger}$ | N/A | N/A | Reversal time | 2cf, S2d |
| Convex policy with gain (without IOFC) | $10^{-2.5}$ | $10^{0.75}$ | $10^{-1.5}$ | N/A | N/A | Reversal time | 3, S8 |
| Convex policy with gain (with IOFC) | $10^{-2.5\dagger}$ | $10^{1.25}$ | $10^{-1.5\dagger}$ | $10^{-0.75}$ | $10^{1.5}$ | Reversal time | 3, 4, 5, S3, S8, S11, S12 |
| Policy gradient without gain (no noise) | N/A | $10^{-1}$ | N/A | N/A | N/A | Reversal time | S6bc |
| Convex policy without gain (no noise) | $10^{1.5}$ | N/A | $10^{1.5}$ | N/A | N/A | Reversal time | S6bc, S10 |
| Convex policy without gain | M | N/A | M | N/A | N/A | Learning perf. | S6def |
| Convex policy with gain (without IOFC) | M | M | M | N/A | N/A | Learning perf. | 2b, S6f |
| Convex policy with gain (without IOFC) | $10^{-1.5}$ | $10^{1.25}$ | $10^{1.5}$ | N/A | N/A | Learning time | S7, S8 |

N/A stands for no value available because the parameter is not applicable for the model in question.
M stands for multiple values available, because the optimization was run multiple times (e.g. for different numbers of distractors).
† means that the parameter value was copied from another parameter optimization procedure.
\* The column for $\eta^V$ was used to indicate the Rescorla-Wagner learning rate $\eta^{\text{RW}}$ , see equation (25).

### Model evaluation

In order to evaluate the computational model simulations and compare them to experimental observations, we employed different metrics:

#### Performance

The learning progress was quantified by tracking the performance during the simulation. The performance is computed as a moving average of the number of correct actions taken within a window of 100 trials. This was implemented with a convolution using a reflection padding of 50 trials at the boundaries (beginning / end of training or before / after reversals). The performance threshold to achieve expertise was set at 85%.

#### Relative gain

The relative gain of the sensory neuron *i* used in Fig. 4a quantifies how the gain 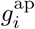 of a pyramidal neuron *i* compares to the gain of the other neurons. It is defined as the gain divided by the average gain across all other sensory neurons,

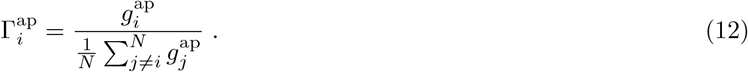

The relative gain for the go-stimulus selective neuron 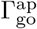 will be taken for a different sensory neuron depending on the current task rule (as in Extended data Fig. S11a), unless it is specified for an explicit rule (as in Fig. 4)c).

#### *π*-weight

The strength of synaptic weight 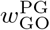 originates from the go-stimulus neuron (the s_1_-selective neuron during the original rule and the s_2_-selective neuron in case of the reversed rule) and targets the go action neuron in the policy network *π*^PG^. The *π*-weight measures 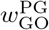 and characterizes the sensorimotor association that has to be learned for the go/no-go sensory discrimination task.

#### Signal-to-noise ratio

The signal-to-noise ratio (SNR) used in Fig. S5 is defined as the ratio between the variance of a stimulus encoding neuron and of the distractor neurons. Since the activity *z*_*s*_ of a go/no-go stimulus encoding neuron corresponds to a Bernoulli variable (with probability *p* = 0.5) amplified by a gain *g*_*s*_ and the sum of distractor neuron activities 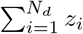) corresponds to a binomial distribution (also with probability *p* = 0.5 and in this example without gain modulation), we find that

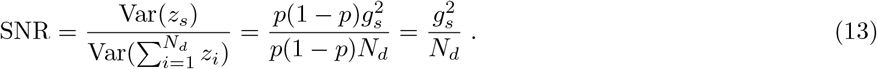

### Animals

C57BL/6, male mice aged between 8 and 16 weeks were used in our experiments. All experimental procedures were conducted in accordance with the requirements of the United Kingdom Animals (Scientific Procedures) Act 1986 and the Federal Veterinary Office of Switzerland, approved by the Cantonal Veterinary Office in Zürich. Experiments were conducted under the authority of Home Office Project License PABAD450E and personal license numbers 285/2014 and 234/2018. Mice were housed at 24°C in a 12-h reverse dark-light cycle (07h00 to 19h00). At the end of the experiment, the mice were deeply anaesthetized, transcardially perfused and euthanized by exposure to CO2 in their home cage.

### Water scheduling

During behavioral training, mice stayed on a 5/2 days water restriction paradigm. They were weighed daily and kept at 85-90% of the previously determined baseline weight. In case animals did not receive enough reward during training sessions (40ml/kg/day), they were given water after training. Water was always provided ad libitum during weekends.

### Handling and habituation

To familiarize the animals with the experimenter, handling was started before surgeries and continued during the recovery period. Habituation to head restraint in progressively lengthened sessions until they tolerated at least 15 minutes of head restraint.

### Cranial window and cannula implantation

Mice were implanted with a cranial window (S1) or cannula (lOFC) under controlled anaesthesia [19]. For the cranial window preparation, a 4 mm cover glass was placed inside the craniotomy and the edges were sealed with dental cement. For imaging deeper structures, the superficial brain tissue was aspirated, a cannula with a glass bottom placed on top of the target area and the edges sealed with dental cement. A head post was attached using dental cement, and the skin was reattached to the implant.

### Sensory learning and reversal learning task

We trained mice on a tactile Go/No-Go reversal learning task [19]. Animals were presented with two sandpaper textures (P100, coarse, s_1_; P1200, fine, s_2_) and they needed to learn to discriminate to either earn a sucrose water reward (‘Go’-texture, P100, ‘hit’) or withhold a response to avoid punishment (‘No-Go’-texture, P1200, ‘CR’, correct rejection). Responding towards the ‘No-Go’-texture resulted in experiencing white noise (‘FA’, false alarm). After animals displayed stable expert performance (d’*>*1.5 in three consecutive sessions; d’= *Z*(hit*/*(hit+miss))−*Z*(FA*/*(FA+CR)), with *Z*(*p*), *p* ∈ [0, 1], being the inverse of the cumulative Gaussian distribution), stimulus-outcome associations were reversed (P100, ‘No-Go’-texture; P1200, ‘Go’-texture) and mice needed to adapt their responses accordingly. The task was amended into a serial reversal learning task by introducing a second rule-switch after animals displayed stable expert performance during initial reversal learning (d’*>*1.5 in three consecutive sessions), reinstating initial stimulus-outcome (P100 as ‘Go’-texture, P1200 as ‘No-Go’-texture).

### Two-photon Ca^2+^ imaging

Neurons were imaged using a custom-built two-photon microscope equipped with a Ti: Sapphire laser system (approximately 100 femtosecond laser pulses; Mai Tai HP, Newport Spectra Physics), a water-immersion 16X Olympus objective (340LUMPlanFl/IR, 0.8 numerical aperture, NA), galvanometric scan mirrors (model 6210; Cambridge Technology) and a Pockels Cell (Conoptics) for laser intensity modulation. Virally expressed GCaMP6f in L2/3 neurons in S1 were imaged at 940 nm. Neuronal activity was recorded at a 15 Hz frame rate (796 × 512 pixels) for 10 seconds each trial by detecting fluorescence changes using a photomultiplier tube (Hamamatsu). Imaging was intermitted during the 3s ITI.

### Ca^2+^ imaging analysis

Ca^2+^ imaging data of each session was processed using the Python-based Suite2p processing pipeline [90]. The recordings were motion-corrected using non-rigid registration. Cells were detected with automated ROI detection followed by manual curation. Raw fluorescence time courses were extracted as each ROI’s (non-weighted) mean pixel value. In addition, a neuropil trace was computed for each ROI to generate neuropil-corrected calcium traces F(t). The fluorescence change during each trial epoch (stimulus-evoked, reward-responsive responses) was then calculated by subtracting the baseline fluorescence F_0_ (mean fluorescence value of the first 1.5 s) from the corrected traces F and dividing by F_0_; Δ*F/F*_0_ = (*F* − *F*_0_)*/F*_0_.

### Chemogenetic silencing

Inhibitory DREADDs (hM4Di) were either expressed in excitatory (CaMKII*α*-hM4D(Gi)-mCherry) or GABAergic VIP-interneurons (AAV1/2-hSyn1-dlox-hM4D(Gi)-mCherry(rev)-dlox, VIP-cre mouse line) in layer 2/3 of the lOFC through virally mediated transfection. hM4Di was activated via intraperitoneal (i. p.) injection of the ligand clozapine N-oxide (CNO dihydrochloride, 1–5 mg/kg, Tocris Bioscience, Catalog number 4936) 25-30 minutes before each training session.

## Data availability

The experimental data that support the finding of this study are available upon reasonable request from the corresponding author. The simulated data can be reproduced from the code provided.

## Code availability

The code for the simulations, statistical analysis and visualizations is provided on https://github.com/unibe-cns/TopDownOFC. The code for data processing of the experimental data is uploaded to https://github.com/banerjeelaboxford.

## Acknowledgements

This works was supported by the Swiss National Science Foundation (project grants 170269 and 192617 to FH, Sinergia grant 180316 to FH and WS), the European Union’s Horizon 2020 research and innovation programme through the ICEI project under the grant agreement No. 800858 (Human Brain Project, to WS), the European Research Council (ERC Advanced Grant BRAINCOMPATH, project 670757, to FH), a Wellcome Trust institutional strategic award (to AB), a Royal Society research grant (RGS\R2\202155, to AB), and a Wellcome Trust Career Development Award (to AB). AB is affiliated to the Digital Environment Research Institute at QMUL. The authors also wish to thank Johanni Brea and Alireza Modirshanechi for helpful discussions.

## Author contributions

MT, AB and WS conceptualized and wrote the paper. Experimental investigations were undertaken by JT and supervised by AB. Computational and mathematical investigations were carried out by MT and supervised by WW and WS. Visualization was performed by MT, JT, AB and WS. Funding was acquired by WW, FH, AB and WS. All authors contributed to reviewing and editing.

---

**Fig. S1.**
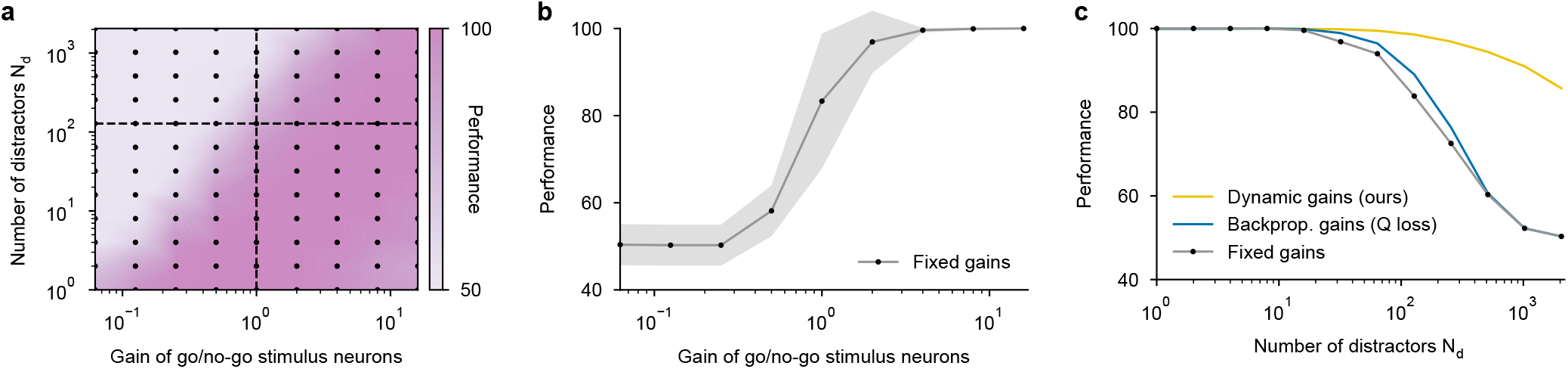
Gain adaptation supports Q-learning from large sensory representations. Same panels as Fig. 1e, f, h using a softmax policy based on estimated Q-values instead of a network trained with policy gradient. **a**, Performance without gain adaptation on a go/no-go sensory discrimination task depending on the number of distractor neurons and on the gain of the two sensory neurons selective for either s1 or s2 (Fig. 1c). The gain of the distractor neurons was always set to 1. Performance was evaluated after 600 trials of training. Each black dot corresponds to the average from 64 different pseudorandom initializations. **b**, Horizontal slice of panel a with 128 distractor neurons and different fixed gains for the go and no-go stimulus encoding neurons. The gray area shows the standard deviation. **c**, Performance achieved with and without dynamic gain modulation, depending on the number of distractor neurons. Gain adaptation was learned either directly through reward-association (yellow) as described in equation (5) or by backpropagating the Q-learning gradient to update a multiplicative gain parameter (blue), see equation 24. The values without gain adaptation (gray) correspond to the vertical slice in panel a.

**Fig. S2.**
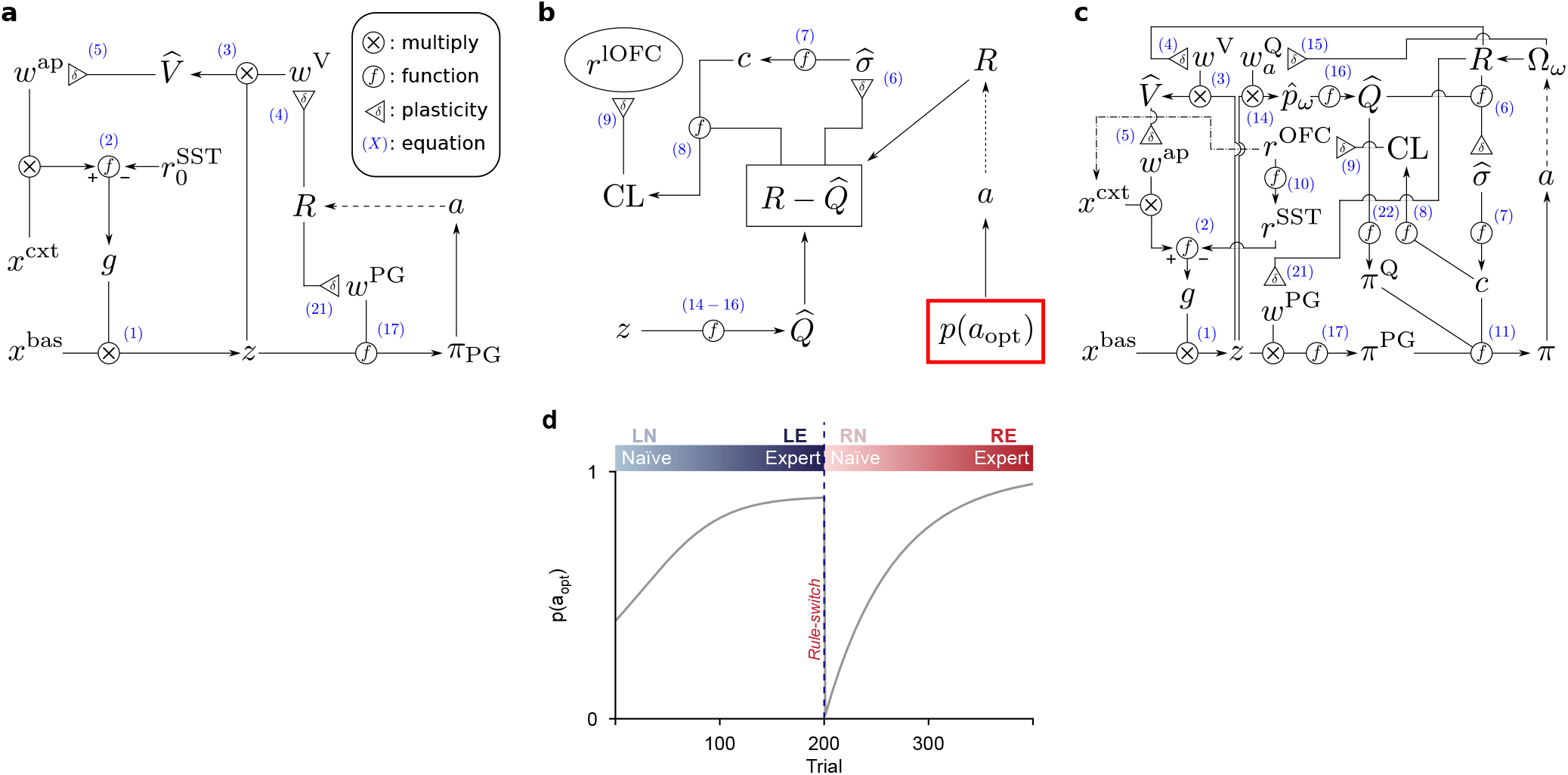
Network map of the model and its equations. **a**, Map of the gain modulation apparatus without the lOFC and using pure policy gradient for action selection. This model was used in the simulations for Fig. 1 to demonstrate, how apical gain amplification based on value facilitates online policy gradient learning from sensory representations encoding distracting information. The numbers in blue refer to the equation numbers for mathematical descriptions. **b**, Network describing how the lOFC activity was modeled for the simulations in Fig. 2, at which point the top-down influence of the lOFC to the sensory representation *z* was not yet included. In order to nonetheless use a realistic evolution of the action selection performance, the policy was replaced by a fixed trace dictating the evolution of the correct action probability *p*(*a*_opt_). **c**, Network schematic of the full model containing the two suggested top-down influences of the lOFC onto the sensory representation (one via SST interneurons and one switching the context representation ***x***^cxt^). Variations of the full model are described in Supplementary Methods 3 **d**, Fixed evolution of the correct action probability used in the simulations described in panel b.

**Fig. S3.**
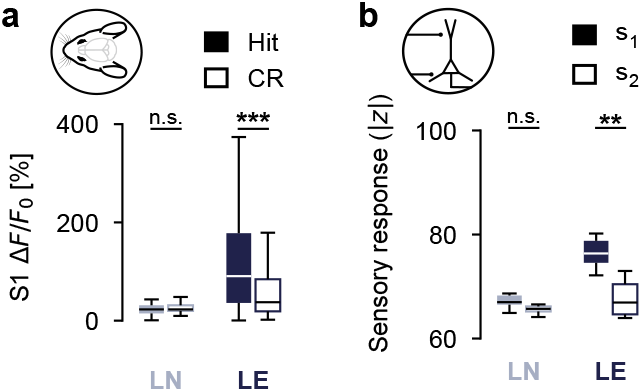
Reward association amplifies stimulus response. **a**, Calcium imaging of mouse S1 during the stimulus presentation window (50, 50 neurons from 4 mice for LN, LE respectively), either before (LN) or after acquisition (LE) of a go/no-go texture discrimination task. Hit trials occur when the reward associated stimulus (s_1_) was correctly responded to with the go action. CR trials occur when the unrewarded stimulus (s_2_) was correctly responded to with the no-go action. Statistical differences were tested by two-sided Welch’s t-test with Bonferroni correction; LN (t(71)=-0.67, p_adj_=1.0; Δ=-3.8, 95% CI [-14.9, 7.4]), LE (t(67)=3.6, p_adj_=0.0011; Δ=71, 95% CI [32, 111]). **b**, Summed firing rates of the simulated sensory neurons, ∑_*i*_ *z*_*i*_, depending on the sensory stimulus. Stimulus 1 (s_1_) is associated with reward and stimulus 2 (s_2_) is not; LN (t(15)=2.5, p_adj_=0.051; Δ=1.8, 95% CI [0.26, 3.4]), LE (t(17)=3.2, p_adj_=0.0096; Δ=8.4, 95% CI [2.9, 13.9]).

**Fig. S4.**
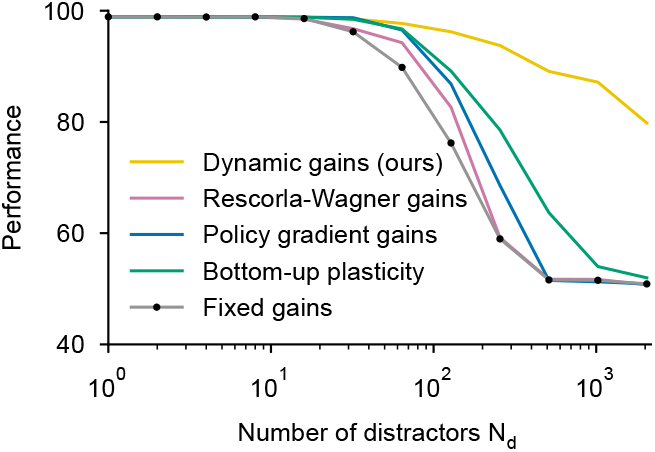
Comparison of robustness to distracting inputs between alternative strategies. Same as Fig. 1h with additional algorithms added for comparison. All variations use policy gradient to learn the synaptic weights of the action selection policy. Yellow: Gain adaptation model based on reward-association, see equation 5. Purple: Gain adaptation according to Rescorla-Wagner. Blue: Adaptation of multiplicative gain parameters according to the backpropagated policy gradient loss. Green: Bottom-up synaptic plasticity dictated by the backpropagated policy gradient. Gray: No gain adaptation in the sensory representation. Each of the alternative approaches are described in more details in the Supplementary Methods 4.

**Fig. S5.**
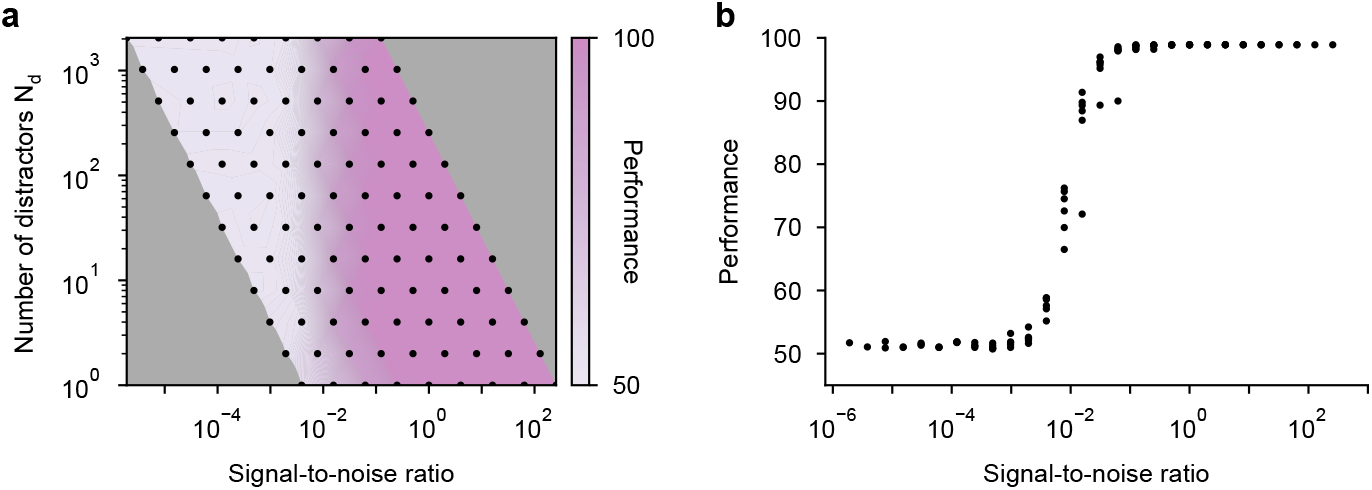
Signal-to-noise ratio determines learning. **a**, Final performance for the model without gain modulation (same as Fig. 1e) depending on the number of distracting neurons (N_d_) and the signal-to-noise ratio (SNR), see equation (13). **b**, From panel a, we suspect that the signal-to-noise ration (SNR) makes a good prediction for the final performance of the model without gain modulation. This can be emphasized by plotting the performances, but only with regard to the signal-to-noise ratio. Each point corresponds to a point from panel a.

**Fig. S6.**
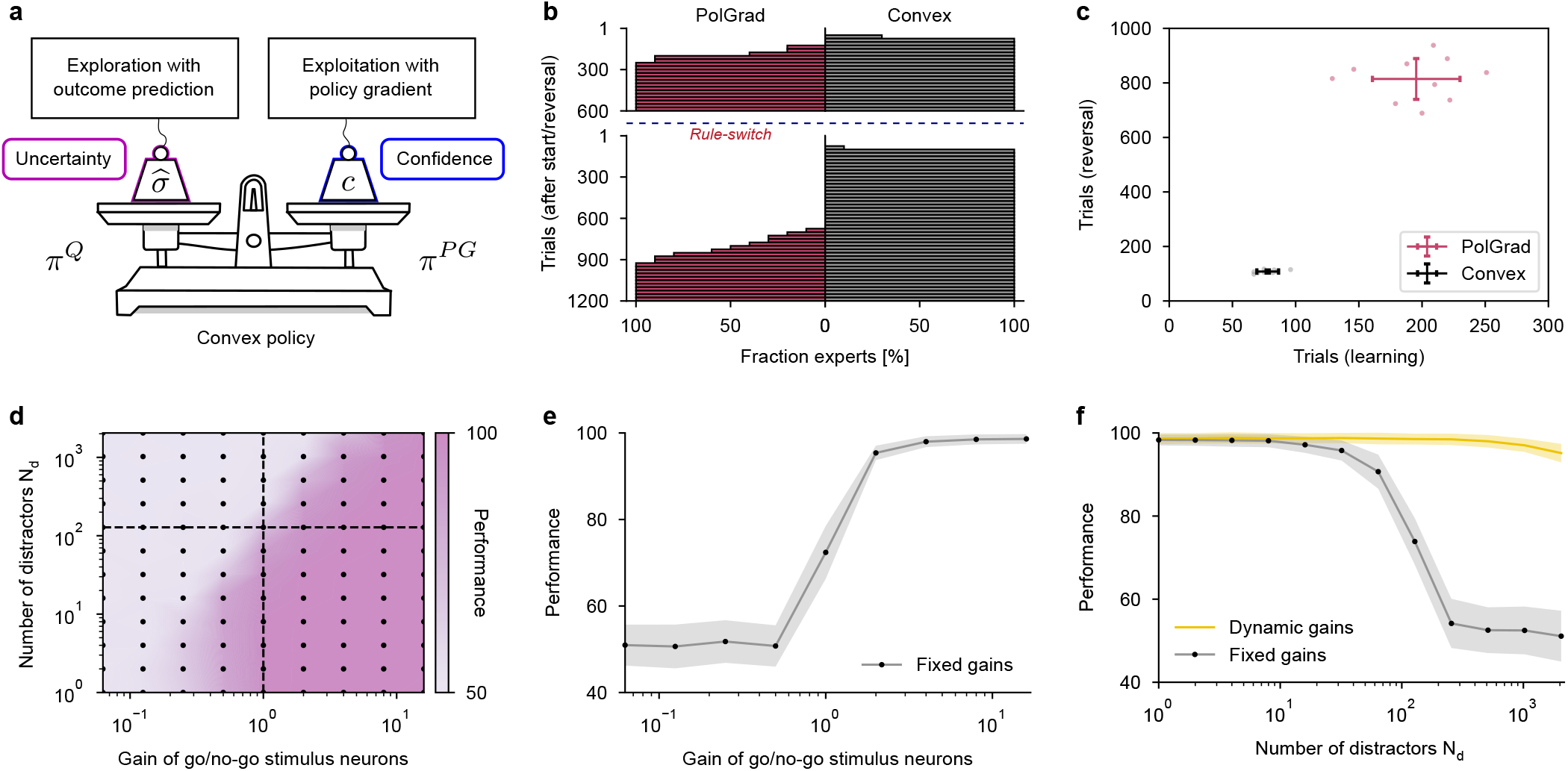
Convex policy for action selection in non-stationary environment. **a**, Schematic of the convex action selection policy, which weighs between an exploitation policy and an exploration policy according to the prediction confidence. **b**, Cumulative distributions of number of trials necessary to reach expert performance for either the original and the reversed rule. This evolution is plotted for the policy gradient method (PolGrad) and for the convex policy (Convex) in both cases without any distractors. In both cases the parameters were optimized for the final performance after the rule reversal. **c**, Number of trials to reach expertise (before and after reversal in the x-axis and y-axis respectively) for the policy gradient and for the convex method. The convex method adapts more rapidly and more consistently. **d, e** and **f** show the same as Fig. 1b, 1c and 1f for a convex action selection policy instead of pure policy gradient. This confirms that the dependence on a high signal-to-noise ratio and the advantage of task-dependent gain modulation also applies for the convex policy.

**Fig. S7.**
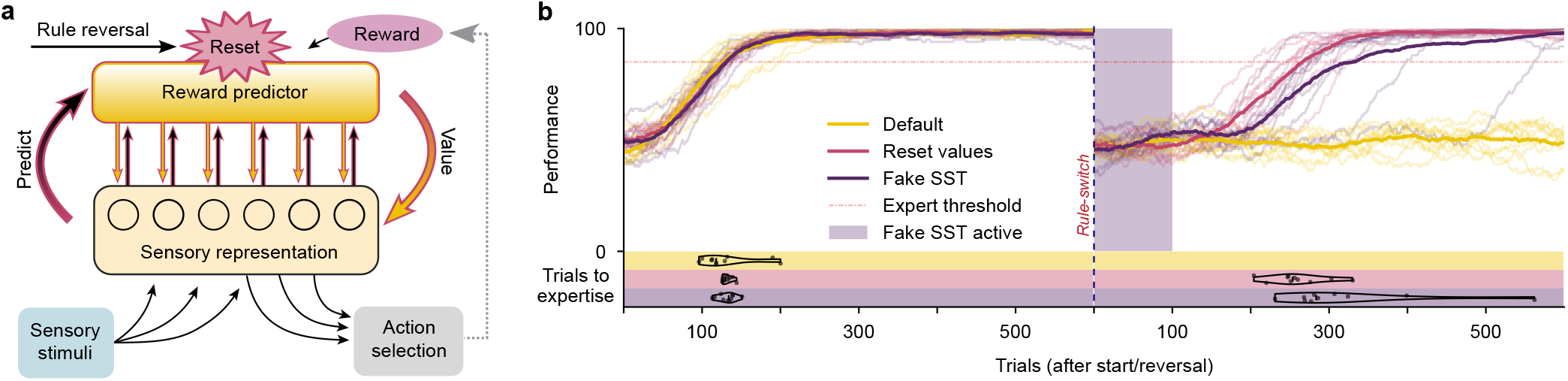
Artificial reset of top-down modulation upon rule reversal. **a**, The ‘Reset values’ model helps us study the importance of top-down gain adaptation during reversal learning by artificially resetting the apical modulation apparatus after a rule change (see Fig. 1e for comparison). Upon rule reversal, the reward predictor and the top-down apical synapses are reinitialized and restart learning from scratch, which negates any bias produced by learning from the previous rule. **b**, Comparing the performance traces of different gain adaptation algorithms shows that apical dendrite inhibition after a rule reversal can help unbias the sensory representation and support reversal learning. The default gain modulation model without lOFC (yellow) struggles to learn after reversal. The ‘Reset values’ model (red) performs much better, since it does not suffer from ill-suited top-down amplification after the reversal. Finally, the ‘Fake SST’ simulation (violet) was tested, which took the default model without lOFC and increased the SST interneuron activity to a high value (10) for a duration of 100 trials after the rule reversal. The fact that the ‘Fake SST’ model adapted almost as well as the artifical ‘Reset values’ model shows that well timed apical top-down inhibition via SST interneurons can provide a biological mechanism to unbias the sensory top-down amplification.

**Fig. S8.**
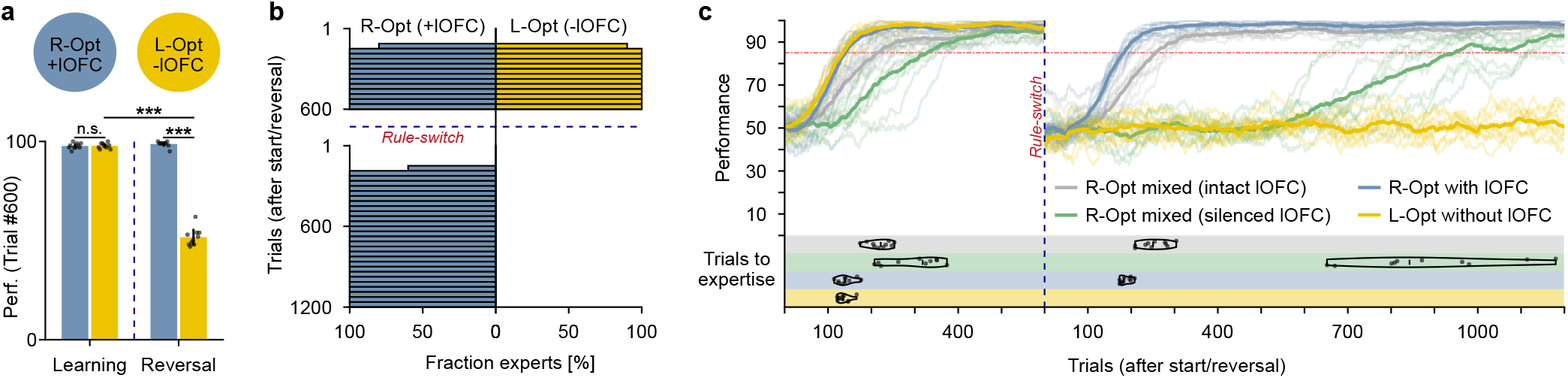
Alternative parameter optimization. **a**, The main model parameters (used in the main figures) are derived to be the same in the simulations where the lOFC is intact or silenced (see Supplementary Methods 5). For completeness purposes, here we also show the performance of the model without lOFC (-lOFC), where the parameters are optimized for performance in the original rule (L-Opt). These are compared to the model with lOFC (+lOFC), where all parameters are optimized for performance in the reversed rule (R-Opt); Learning +lOFC vs. -lOFC (t(18)=-0.16, p_adj_=1.0; Δ=-0.10, 95% CI [-1.4, 1.2]), Reversal +lOFC vs. -lOFC (t(11)=32, p_adj_*<*0.001; Δ=47, 95% CI [44, 50]), Learning vs. Reversal for -lOFC (t(11)=31, p_adj_*<*0.001; Δ=46, 95% CI [43, 49]). **b**, The cumulative distribution of expert agents depending on the trial before and after reversal, again for the model without lOFC optimized for the first learning phase, L-Opt (-lOFC) and the same for the model with lOFC optimized for the reversal phase, R-Opt (+lOFC). **c**, Comparison of the performance traces the model without lOFC optimized for the learning phase (L-Opt without lOFC, yellow) and the model with lOFC optimized for the reversal phase (R-Opt with lOFC, blue) with the model optimized jointly for the reversal phase (R-Opt mixed) either with the lOFC intact (green) or silenced (gray).

**Fig. S9.**
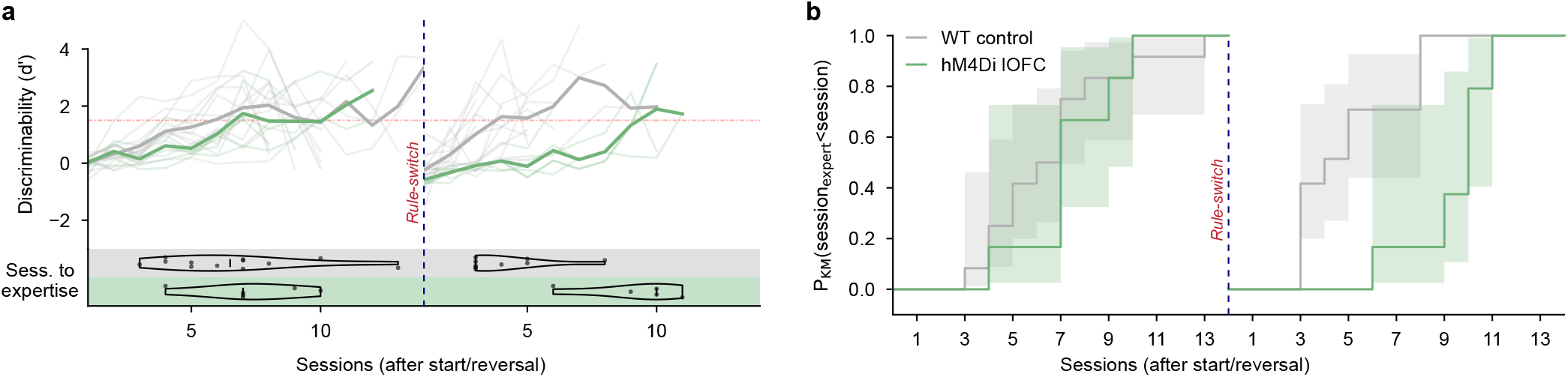
Lateral OFC inhibition impairs reversal learning. **a**, Performance of wild-type (WT, gray) and lOFC silenced (green) mice during reversal learning (12 and 6 mice, respectively). Bottom: violin plots show the distributions of the number of sessions necessary to become expert for the original or the reversed rule. **b**, Kaplan–Meier curves for the number of sessions to reach expert performance. Expertise was defined as the first session at which performance met or exceeded the expert threshold. Animals that had to be aborted from training before reaching expertise were treated as right-censored at their last session. Statistical differences were assessed using a two-sided log-rank test with Bonferroni correction and effect size was estimated with a Cox proportional hazards model; Learning (Log-rank: *χ*^2^(df=1)=0.23, p_adj_=1.0; Cox PH: HR=1.18, 95% CI [0.44, 3.21]), Reversal (Log-rank: *χ*^2^(df=1)=8.4, p_adj_=0.0076; Cox PH: HR=12.0, 95% CI [1.6, 91.0]).

**Fig. S10.**
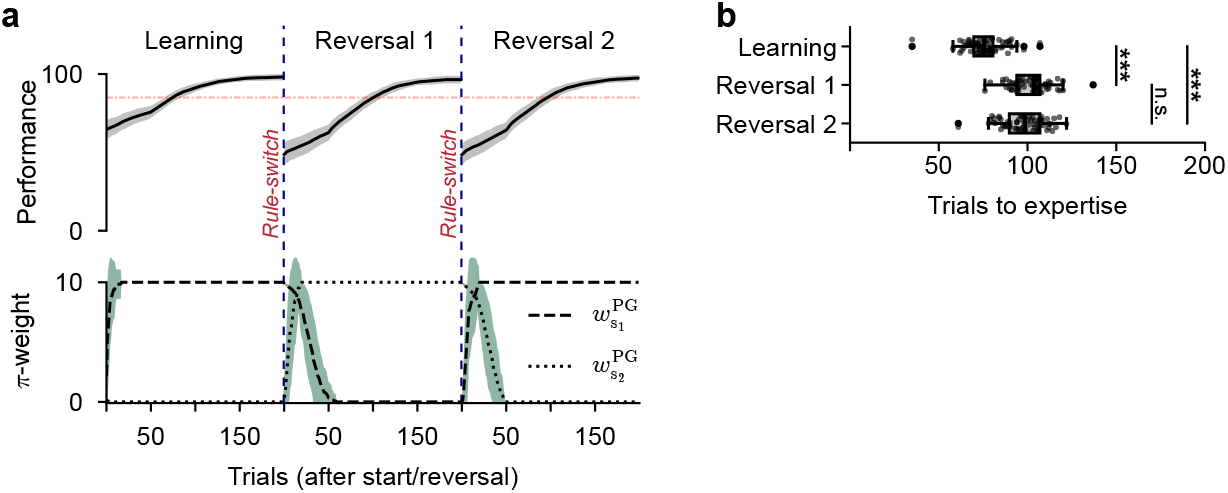
Catastrophic forgetting without gain modulation. **a**, Evolution of the performance trace (top) and *π*-weights (bottom) without gain modulation and without distracting stimuli. The policy weights are unlearned after each reversal in order to allow the behavior to adapt to the new rule. **b**, Number of trials necessary to reach expert performance (85%) during the first learning phase (Learning), after one rule reversal (Reversal 1) and after the second reversal (Reversal 2). Statistical differences were tested using a two-sided log-rank test with Bonferroni correction and effect size was estimated with a Cox proportional hazards model; Learning vs. Reversal 1 (Log-rank: *χ*^2^(df=1)=107, p_adj_*<*0.001; Cox PH: HR=8.2, 95% CI [5.3, 12.7]), Learning vs. Reversal 2 (Log-rank: *χ*^2^(df=1)=84, p_adj_*<*0.001; Cox PH: HR=6.3, 95% CI [4.1, 9.6]), Reversal 1 vs. Reversal 2 (Log-rank: *χ*^2^(df=1)=1.33, p_adj_=0.75; Cox PH: HR=0.83, 95% CI [0.59, 1.17]).

**Fig. S11.**
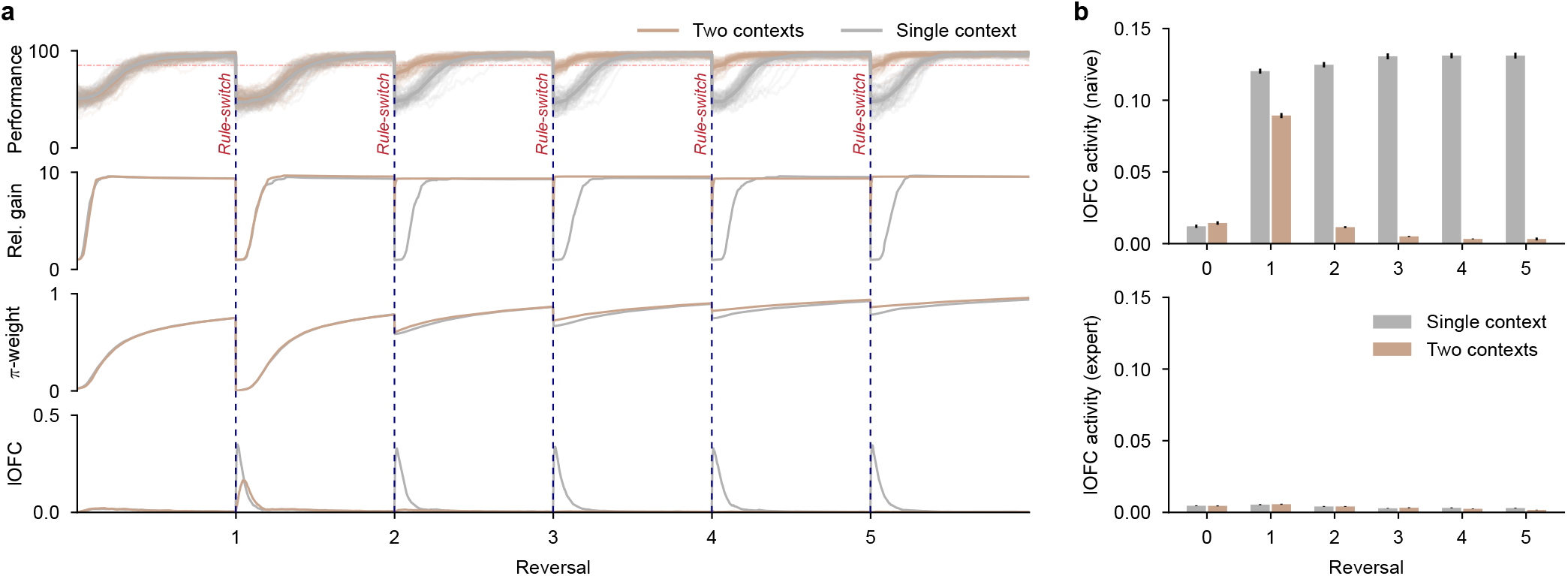
lOFC as context switch. **a**, Average performance trace during five sequential rule reversals (top row) as well as the evolution of the average relative gain and the average *π*-weight for the go-stimulus neuron of the current rule (middle rows). The rapid adaptation of the relative gain to the go-stimulus explains the increased adaptation speed shown in Fig. 4g. It also shows the evolution of the lOFC activity (bottom) modeled by the context-prediction error (CPE). All the mean traces are shown both for the model with a single context representation (gray) and with two context representations (brown). **b**, Average lOFC activity (modeled as CPE) either during the naïve phase (first 100 trials) or the expert phase (last 100 trials) of each reversal cycle (of 600 trials each). The error bars show the standard error.

**Fig. 12.**
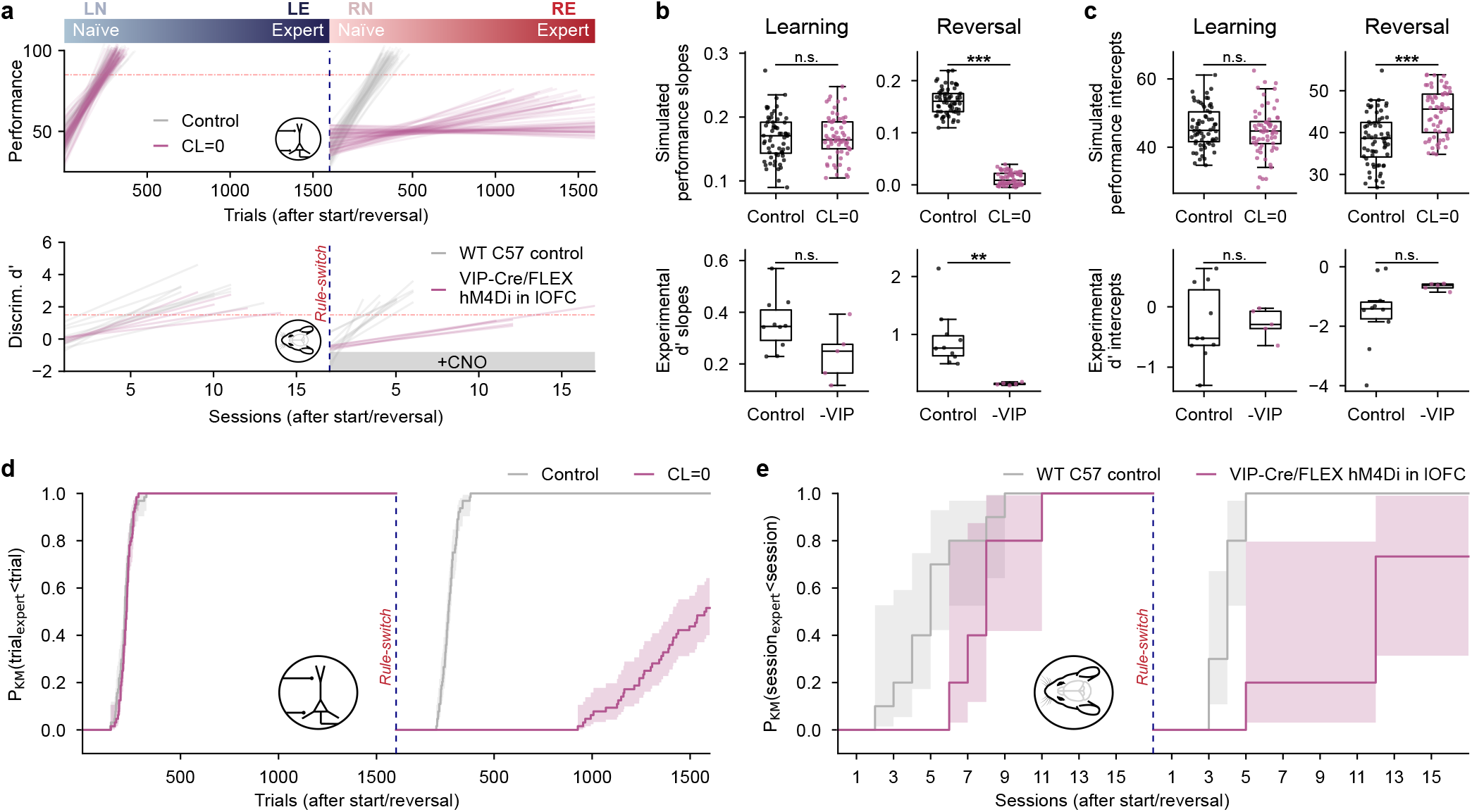
lOFC VIP interneurons support reversal learning. **a**, Linear regressions of the behavioral performance traces of both model (top) and mice (bottom) during reversal learning shown in Fig. 5a. **b**, Distributions of the slopes of the linear regressions before (left) and after reversal (right) compared between the control and the lOFC VIP interneuron inhibited condition for simulations (top) and mice (bottom); simulations during Learning (t(126)=-0.032, p_adj_=1.0; Δ=-0.0002, 95% CI [-0.012, 0.012]), simulations during Reversal (t(91)=43, p_adj_*<*0.001; Δ=0.15, 95% CI [0.14, 0.16]), mice during Learning (t(7.9)=2.0, p_adj_=0.33; Δ=0.12, 95% CI [-0.02, 0.25]), mice during Reversal (t(9.1)=4.9, p_adj_=0.0031; Δ=0.77, 95% CI [0.42, 1.1]). **c**, Distributions of the intercepts of the linear regressions before (left) and after reversal (right) compared between the control and the lOFC VIP interneuron inhibited condition; simulations during Learning (t(123)=1.0, p_adj_=1.0; Δ=1.2, 95% CI [-1.1, 3.4]), simulations during Reversal (t(125)=-6.1, p_adj_*<*0.001; Δ=-6.2, 95% CI [-8.3, -4.2]), mice during Learning (t(13)=-0.090, p_adj_=1.0; Δ=-0.02, 95% CI [-0.51, 0.47]), mice during Reversal (t(9.4)=-2.4, p_adj_=0.16; Δ=-0.89, 95% CI [-1.72, -0.05]). **d**, Kaplan–Meier curves (see Fig. S9b) for the number of sessions the simulated agents needed to reach expert performance. Statistical differences were assessed using a two-sided log-rank test with Bonferroni correction and effect size was estimated with a Cox proportional hazards model; Learning (Log-rank: *χ*^2^(df=1)=0.49, p_adj_=0.97; Cox PH: HR=1.13, 95% CI [0.79, 1.60]), Reversal (Log-rank: *χ*^2^(df=1)=155, p_adj_*<*0.001; Cox PH: HR=165, 95% CI [39, 707]). **e**, Same as d for mice; Learning (Log-rank: *χ*^2^(df=1)=3.4, p_adj_=0.13; Cox PH: HR=2.7, 95% CI [0.84, 8.7]), Reversal (Log-rank: *χ*^2^(df=1)=10.2, p_adj_=0.0028; Cox PH: HR=14.7, 95% CI [1.97, 109.98]).

## Algorithm 1 Simulation of one trial

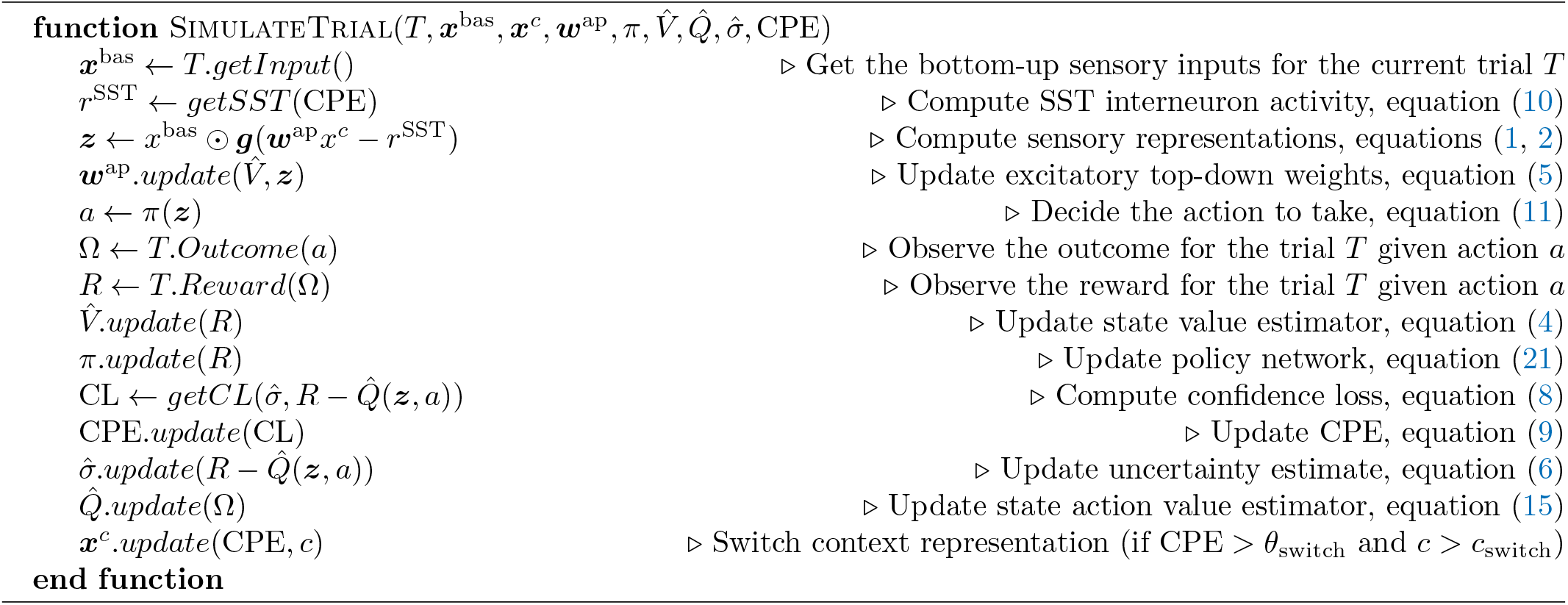

## Algorithm 2 Iterative grid search parameter optimization

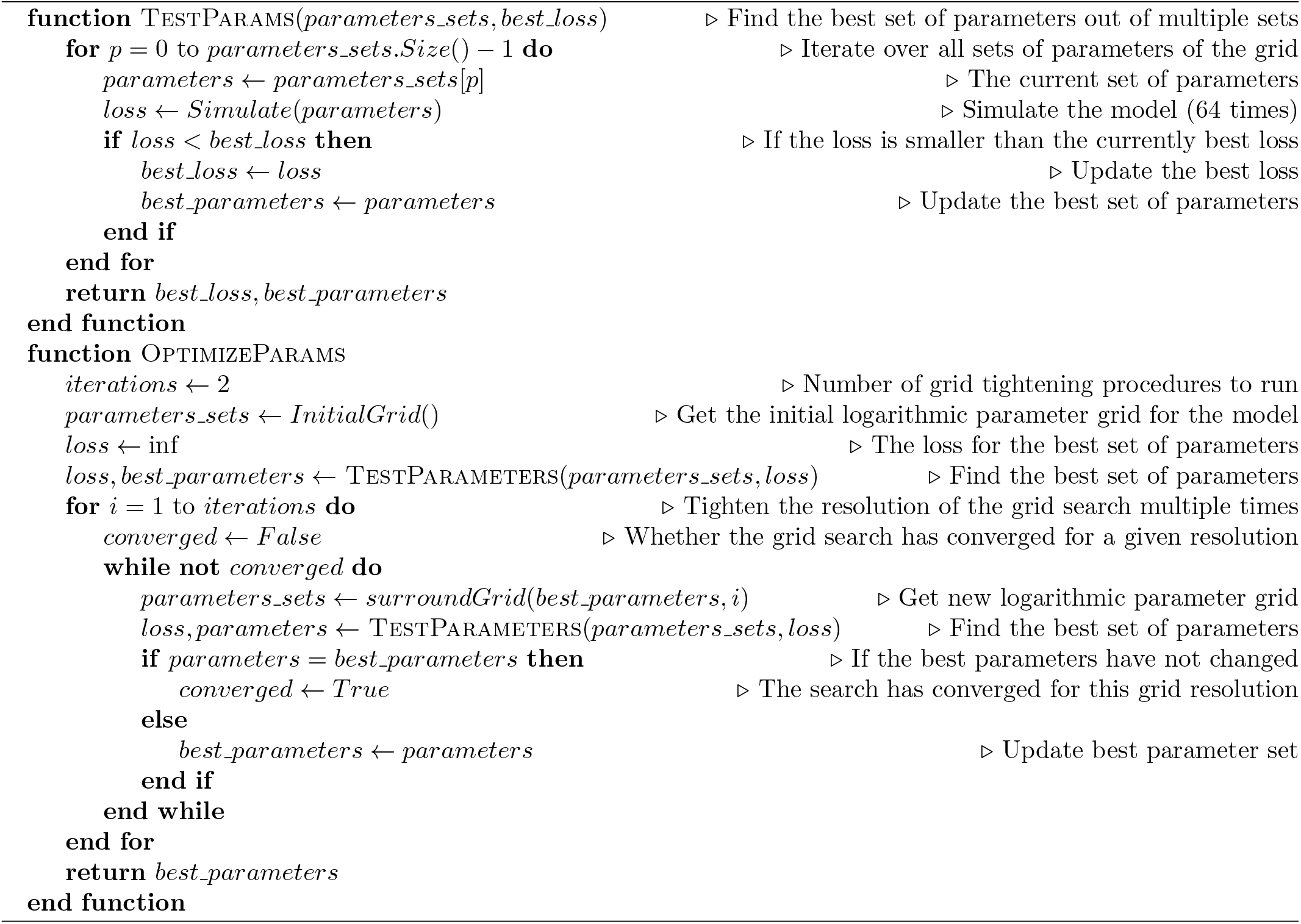

## Supplementary Methods

### 1 Inferring action-values based on the outcome prediction network

The action-value estimate 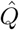 was computed by using an outcome prediction network. For each outcome *ω* ∈ {reward, punishment}, we defined the indicator function Ω_*ω*_, which indicated whether an outcome *ω* took place (Ω_*ω*_ = 1) or not (Ω_*ω*_ = 0). The estimated probability to encounter outcome *ω* upon choosing action *a* ∈ {go, no − go} in response to stimulus representation ***z*** is denoted by 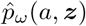. We modeled these estimated outcome probabilities by means of 4 perceptrons, one for each of the 4 action-outcome pairs (*a, ω*), with a standard logistic activation function *φ*,

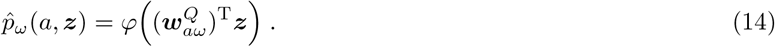

These probability estimators were trained online, each time an action *a*^*′*^ was taken according to

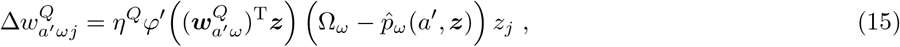

where *φ*^*′*^ denotes the derivative of *φ*. Because the state-action-outcome triples (***z***, *a, ω*) define the rule, the outcome predictions 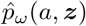 represent the internal understanding about the rule (and more generally about the context). With these outcome predictors, 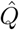 can be expressed by a sum weighted according to the value *R*_*ω*_ ∈ {−1, 1} of each outcome

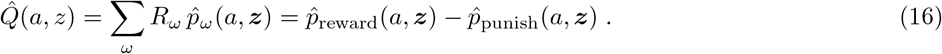

We set two possible outcomes, *R*_reward_ = 1 and *R*_punish_ = −1, but the model can also be generalized to more outcome types.

### 2 Action selection by policy gradient and action-value estimate

Action selection was simulated to be based on the sensory representation ***z***. The policy network consisted of a perceptron with sigmoidal transfer function *ϕ*, which produced the action selection probability *π*^PG^,

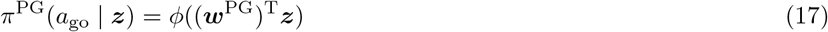

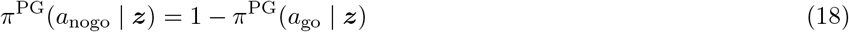

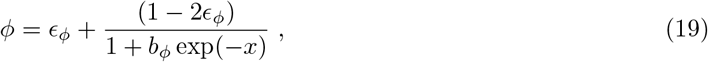

where *ϵ*_*ϕ*_ and *b*_*ϕ*_ are constants. The network learned through policy gradient [85] by minimizing the loss

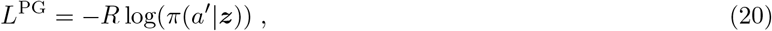

where *a*^*′*^ is the chosen action and *R* is the reward amplitude of the outcome observed. This loss was minimized by online gradient descent with learning rate *η*^PG^ according to the learning rule

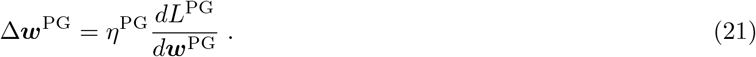

Since a pure policy gradient network approach can lead to an excessively exploitative action selection with very little exploration after convergence, it can be inflexible when facing non-stationary conditions such as rule reversals (Extended Data Fig. S6e). We therefore combined it with a more exploratory policy *π*^*Q*^, which is computed by weighing the predicted outcomes 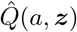, with a softmax transfer function parameterized by a temperature constant *T* ^Q^,

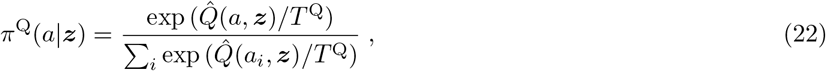

where 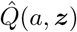 is given by equation (16) and the sum extends across the two actions *a*_*i*_ ∈ *{a*_GO_, *a*_NG_}, corresponding to lick or no lick, respectively go or no-go. Both the exploratory policy *π*^*Q*^ and the exploitative policy *π*^PG^ were incorporated into the final policy *π*, which consisted of the confidence weighted convex combination (Extended Data Fig. S6a),

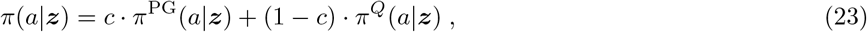

where *c* is the prediction confidence, see equation (7). The overall policy *π* was shown to learn effectively upon reversal learning and therefore ensured that any lack of adaptability would not be caused by the inflexibility in the action selection network (Extended Data Fig. S6e). Depending on the purpose of the modeling, we used different algorithms for the action selection. In most cases, the convex policy was used as described in equation (11). For Fig. 1 however, the purpose was to highlight the impact of the sensory representation *z* and gain modulation for a standard policy, which is why the pure policy network *π*^PG^ was used (for completeness purposes Fig. 1 is replicated with the convex policy *π* in Extended Data Fig. S6). For the first reversal simulations—which illustrated the dynamics of uncertainty, confidence and context-prediction error (Fig. 2b,c,f)—we imposed a performance learning curve, dictated by a fixed evolution of the correct action probability *p*(*a*_opt_) (Extended Data Fig. S2d).

### 3 Model combinations

In this manuscript, we implemented different combinations of action-selection policies and gain modulation mechanisms (Extended Data Fig. S2). The action-selection policy was one of: policy gradient, fixed performance trace (fixed policy), convex policy, predicted reward policy. The gain modulation was one of: none, only reward-guided amplification without lOFC, amplification with lOFC targeting SST interneurons in S1, amplification with lOFC switching the context representation, Rescorla-Wagner, backpropagated gain. Extended Data Table 1 summarizes, which model was shown in which figure. While multiple model combinations were implemented for completeness purposes, three were central to our main findings. 1) To demonstrate the advantage provided by top-down excitation in stationary environments, we combined classical policy gradient for action selection with apical gain amplification of sensory neurons based on reward association (Extended Data Fig. S2a). 2) To illustrate the second-order statistics of reward-prediction errors in a non-stationary environment (without using the full model adapted for the purpose of reversal learning), we used a fixed policy performance trace (Extended Data Fig. S2b). This allowed to generate a realistic evolution of the reward sampling distribution and study the dynamics of confidence loss and context-prediction error during reversal learning (Extended Data Fig. S2d). 3) Finally, the full model with lOFC feedback to S1 SST interneurons and in some instances affecting the context representation used a confidence-weighted convex policy (Extended Data Fig. S2c). The convex policy ensured that any lack of reversal performance would not be due to deficits in the flexibility of the policy (Extended Data Fig. S6), but could be attributed to failing to re-adapt the sensory representation. The pseudocode to simulate one trial of the full model can be found in Algorithm 1. Alternative policies and gain modulation strategies (e.g. expected reward policy, Rescorla-Wagner gain, backpropagated gain) that were only simulated as comparison to the main model are described in the Supplementary Methods 4.

### 4 Model alternatives

To appropriately benchmark and compare the computational model, alternative algorithms and variants of the model were tested (see Extended Data Fig. S1 and S4).

#### Policy based on predicted reward

The reported challenges of the policy gradient algorithm to learn the discrimination task from a sensory representation with many distractors (see Fig. 1e, f) and the increased robustness conferred by the value-based gain modulation (see Fig. 1h) were replicated for a policy based on the reward prediction (see Extended Data Fig. S1). In these control simulations the action selection policy was based on the softmax of the predicted reward conditioned on the action and the state representation with a temperature of 0.1, see equation (22). Using this policy, three variants of neuron-specific gain adaptation in the sensory representation were tested: no gain adaptation, value-based gain adaptation, and backpropagated gradient based gain adaptation. The value-based gain adaptation was the same as in our main model, see equation (5). In the third version, the gain parameter was directly adapted by backpropagating the squared reward prediction error. The resulting gain updates were

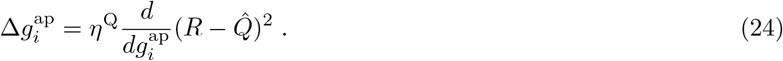

#### Rescorla-Wagner gain

The classical Rescorla-Wagner model [86] describes stimulus-reward association weights 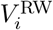 and their updates

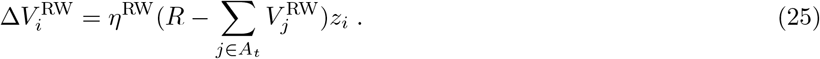

Traditionally, the index *i* would refer to a stimulus *i* and *A*_*t*_ would be the set of stimuli present at time *t*, while *z*_*i*_ would be the salience of stimulus *i*. In our framework adapted for neural sensory representations, however, *i* should be interpreted as the index of a sensory neuron, *A*_*t*_ as the set of active neurons at time *t*, and *z*_*i*_ as the firing rate of neuron *i*. As usual, *R* represents the reward, and *η*^RW^ is the learning rate. In order to test the Rescorla-Wagner rule for gain modulation, the gain of each neuron *i* was modeled as

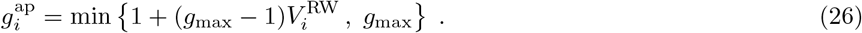

#### Policy gradient gain

As a benchmark, one variant was tested, where the neuron-specific gain parameters of the sensory representation was adapted by the backpropagated policy gradient loss, see equation (20). The resulting updates were

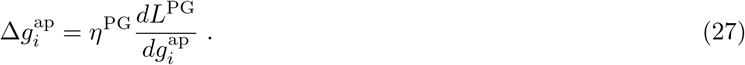

#### Bottom-up plasticity

In the main simulations the basal activation 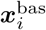 of the sensory neuron *i* was uniquely determined by either the presence of one of the task-relevant stimuli or one of the *N*_*d*_ irrelevant noise inputs (see Simulated go/no-go task). This can be mathematically formalized as

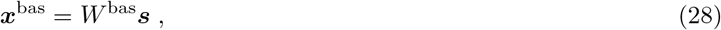

where *W* ^bas^ is the *N × N* identity matrix and ***s*** is an array concatenating the task-relevant stimuli *s*_1_, *s*_2_ with the task-irrelevant inputs {*ξ*_*j*_ |*j* ∈ ℕ, 1 ≤ *j* ≤ *N*_*d*_ }. In order to isolate the effects of top-down adaptation in the simulations, the bottom-up weight matrix *W* ^bas^ was mostly kept fixed. In order to test the effect of bottom-up plasticity, the weights were allowed to adapt following the backpropagated policy gradient loss

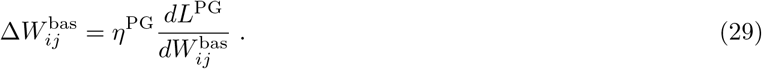

While the main model achieved its performance purely with top-down adaptation and without bottom-up plasticity, the two mechanisms are not mutually exclusive. Hence in principle, the main model is compatible with any bottom-up plasticity rule.

### 5 Parameter values

The parameters 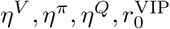 and *w*^SST^ were optimized by grid search, while all remaining parameters had fixed values, which are reported in Extended Data Table 2. The parameter optimization took place by testing different sets of parameters values taken from logarithmic grid. Each set of parameters was tested with 64 pseudo-random initializations. After the best set of parameters was determined, a new tighter logarithmic grid of parameter values was created surrounding the best parameter configurations. Each new set of parameters was then tested to determine, if they could improve on the best configuration. This process was repeated a second time with yet a tighter grid and the final best parameter configurations was chosen, see Extended Data Algorithm 2. The criterion for the best set of parameters was usually the number of trials to reach expert performance after one reversal. In some instances, like in Fig. 1, we optimized for the final performance for the original rule (see Learning perf. in Extended Data Table 3) and in one example, we optimized for the number of trials to reach expert performance in the original rule (Extended Data Fig. S8).

The optimized values are reported in Extended Data Table 3. While many different parameter optimizations were performed, only two are relevant for the main results. 1) For the results in Fig. 1, we optimized the parameters with respect to the final learning performance (without any rule reversal). 2) For the remaining main Figures, we first optimized the model without lOFC to achieve expert performance after one reversal in the least number of trials possible. This allowed as to fix the policy related parameters *η*^*π*^ and *η*^*Q*^, before optimizing the remaining parameters related to the top-down modulation of the sensory representation. While this ‘mixed’ optimization strategy did not target the theoretically best parameter configuration for the full model with lOFC (as illustrated in Extended Data Fig. S8), it prevented any potential bias towards the model with lOFC, when comparing the model with and without lOFC (Fig. 3).

The hyperparameters of the control models (the policy based on Q-values shown in Extended Data Fig. S1, the various alternative sensory representation adaptation strategies shown in Extended Data Fig. S4) were also optimized by the same grid search algorithm. Individual hyperparameter values were optimized for each number of distractors or each fixed gain (as in Fig. 1 or Extended Data Fig. S1, S4, S5, S6). The specific parameter values for all such variations of the task conditions can be found with the code (see Code availability).

## Supplementary Equations

### 1 Gains modulate neuron-specific learning rates

For a rate-based neuron *j*, the multiplicative gain *g*_*j*_ modulates the pre-gain activation *x*_*j*_ to produce the firing rate

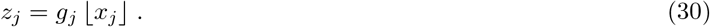

At the same time, most synaptic weight learning rules—from Hebb to error backpropagation—are almost universally proportional to the presynaptic activation. As a consequence, increasing the multiplicative gain of a neuron, raises the rate of change of its efferent synapses. From the perspective of the weight learning rule, the learning rate of a synapse *w*_*ij*_ can be viewed as being modulated by the multiplicative gain *g*_*j*_ of the afferent neuron *j*,

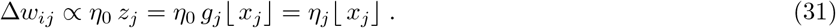

Here, *η*_0_ corresponds to the traditional global learning rate and *η*_*j*_ = *η*_0_ *g*_*j*_ is the learning rate of synapses activated by a neuron *j*. Therefore, by increasing the gain of task-relevant compared to task-irrelevant sensory neurons, it is possible to help the synapses of the policy network to learn more strongly from the task-relevant information. On the other side, reducing the gain will reduce the plasticity. Hence SST interneurons activated by lOFC upon reversal can protect policy weights from being unlearned.

### 2 Apical amplification with respect to task-relevance

In this section, we will elaborate on the model for learning sensory representations that exhibit heightened responses to task-relevant information. For this, we assume that in biological neural networks, the main mechanism to produce selective sensory representations is not based on the synaptic plasticity of feed-forward pathways. Rather, in order to keep a wide range of features available for multiple different contexts, the brain needs to increase the gain of task-relevant neurons via top-down afferents. We further assume that this gain modulation occurs by targeting the apical dendrites of sensory pyramidal neurons. With this in mind, we can set the following objective for the top-down gain modulation mechanism:

#### → High task-relevance of a stimulus should lead to a high gain and low task-relevance a low gain

This objective can be formalized and implemented in different ways. We opted to define the task-relevance of a stimulus *i* in terms of its contribution to the outcome prediction 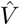,

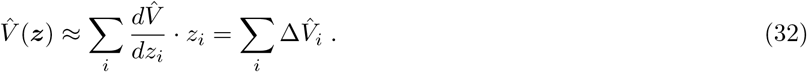

Here the approximation becomes an equality in the case of a linear dependence, which is the case in the model. The term 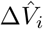 can be interpreted as the reward prediction value gained by the model from the activity of neuron *i*. For apical excitation, we are interested in the the positive part 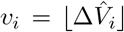, of which the expectation E [*v*_*i*_] makes the definition of task-relevance. In order to produce an apical gain amplification based on the task-relevance of a neuron *i*, we applied the plasticity rule

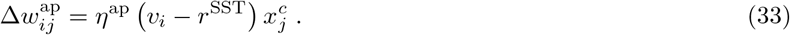

By analyzing this update rule, we can derive the convergence state of the apical excitation for a neuron *i*. For this we will consider the setting, where *j* ∈ {1} and 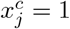. In expectation the synaptic plasticity can hence be formulated as

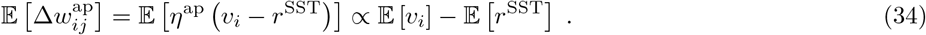

We find that the plasticity rule lets the apical excitation transmitted by the apical synapses 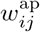 grow large for neurons with a task-relevance greater than the average SST activity, and shrink small otherwise,

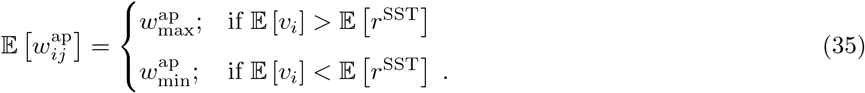

This confirms that 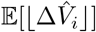 can be considered as out effective definition of task-relevance. The rectifier function ⌊·⌋ is used to prevent the explicit apical inhibition of neurons that are predictive of negative outcomes. Instead, these negative contributions will have the same effect on the gain modulation those of neurons, which are irrelevant for the outcome prediction (i.e. neurons with 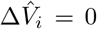). Finally, one can note that the SST activity not only reduces immediate apical activity, but also drives synaptic weight depression in the apical dendrites [87] and thereby raises the task-relevance threshold to drive gain amplification. The SST interneuron activity *r*^SST^ is excited by the lOFC, which increases *r*^SST^ in case of context-prediction error, see equation (10). In consequence, following a rule reversal, the lOFC raises the task-relevance threshold for the consolidation of top-down apical amplification.

### 3 Distinguishing sources of uncertainty

It has been demonstrated that the expected squared error of an estimator can be decomposed into three components: an irreducible, a bias and a variance component [88]. Here, we reproduce this result and adapt it to the case, where distracting information and changes of context come into play. We first define the estimation target *Y* = *f*(*X*) + *ϵ* with E[*ϵ*] = 0 and 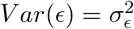. These targets are predicted by a stochastic estimator 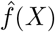 based on the observables *X*. The expected squared prediction error between the target and the estimator can be interpreted as the predictive uncertainty. Next we will demonstrate how the predictive uncertainty can be decomposed into three separate sources of uncertainty: (1) an intrinsic and irreducible error (sometimes referred to as *risk* [89, 90]) determined by *ϵ*, (2) a bias, which is reduced if the expected value of 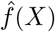 closely matches *f*(*X*) and (3) a variance term which arises from the intrinsic stochasticity of 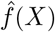:

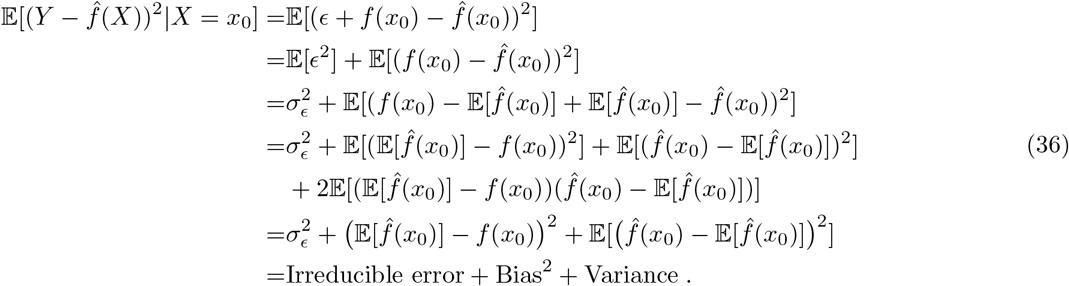

We can also apply this framework to the reinforcement learning model, where the estimation target *Y* represents the reward *R* of the deterministic go/no-go task and the estimator 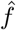 represents the reward prediction 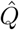. In this case, the observables *X* encodes the action as well as the sensory information. Next we will adapt this decomposition to consider distracting inputs and non-stationary changes of context.

#### Distraction uncertainty

In the simulations, the reward prediction was deterministic and had no intrinsic stochasticity, meaning the variance term in equation (36) was zero. The reward outcomes were also deterministic, meaning the irreducible error 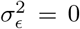. However the predictor also received task-irrelevant information *D*. This contribution can be emphasized by splitting off this distracting information *D* from the observables *X* as distinct inputs to the estimator 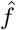. Using the same process as in equation (36), we find

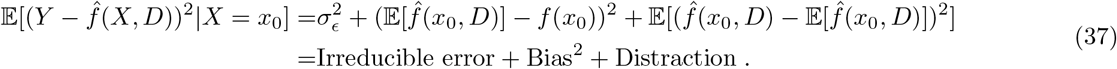

Under this formulation, the top-down amplification of task-relevant signals can be understood as a means to reduce the distraction component of the prediction uncertainty.

#### Contextual uncertainty

So far we assumed a single stationary context. In order to consider the possibility of a change of context, we just need to introduce the dependence of the target *Y* on a current context *c*^∗^, while the model assumes the context *c*^*m*^,

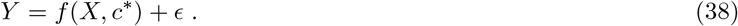

We can now decompose the predictive uncertainty in a non-stationary environment. In case the correct context is assumed (*c*^*m*^ = *c*^∗^),

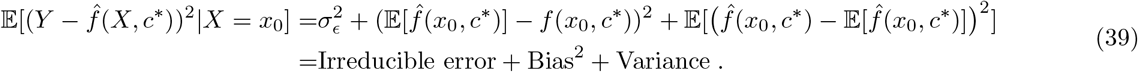

This is similar to what we found in the stationary case, see equation (36). However, if the wrong context is assumed (*c*^*m*^ = *c*^*†*^≠ *c*^∗^), we find that the bias term will be affected by the contextual mismatch

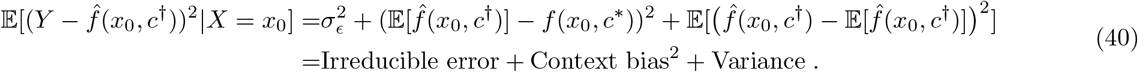

Since the incorrect context assumption would create a strong bias, we refer to this term as context bias to distinguish it from the bias in the stationary case.

#### Unexpected uncertainty

Next we will analyse the dynamics of the prediction uncertainty by looking at the mismatch between the true uncertainty *σ*^2^ and the expected uncertainty 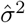 before and after a change of context. So far, to simplify the notation, we only demonstrated how to decompose the conditional expected squared prediction error given the observables *x*_0_. However the analogous decomposition can be made for the full expectation (taken over the intrinsic stochasticity of 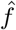 and over *ϵ*, but also over the observables *X*)

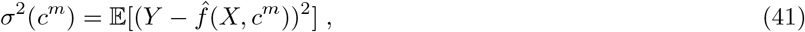

which is the estimation target of the expected prediction uncertainty 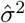, see equation (6). Before a change of context— in case the correct context is assumed (*c*^*m*^ = *c*^∗^) and the uncertainty estimator 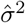 has converged, 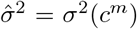—we get

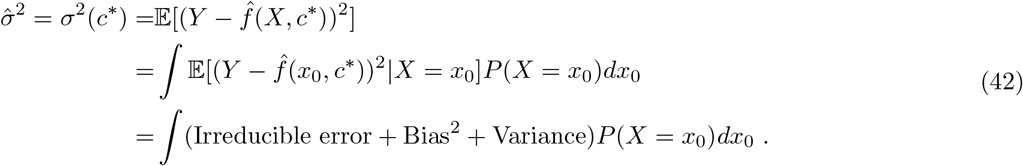

In a scenario, in which learning of 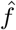 has converged successfully (Bias^2^ = 0), a sudden change of context would increase the squared error by inducing a context bias, see equation (40). This increase in uncertainty can be quantified by the unexpected uncertainty, which is obtained by subtracting the true uncertainty just after a change of context *σ*^2^(*c*^*†*^), with the expected uncertainty just before the change of context 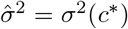, equation (42). Indeed, we find that the unexpected increase in uncertainty, which can be attributed to the change of context, equates to

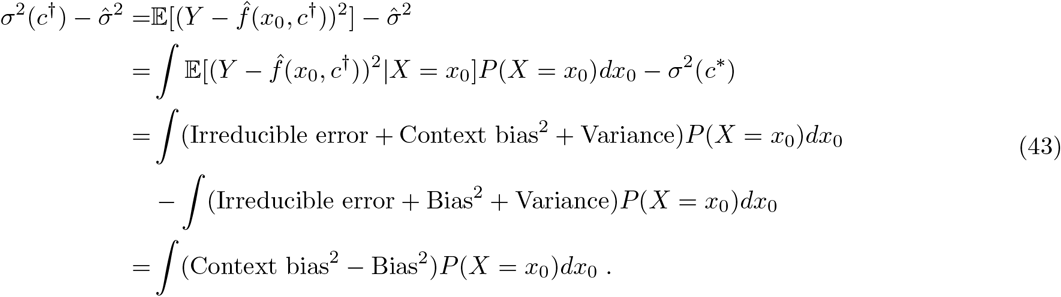

In equation (43), we assumed that the variance term was independent on the assumed context *C*, which is trivial in case the estimator 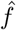 is not stochastic at all. Finally, if learning of 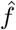 had perfectly converged (Bias^2^ = 0) prior to the change of context,

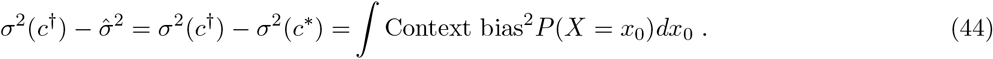

Therefore, in an idealized setting, the unexpected increase in uncertainty following the change of context becomes equal to the contextual uncertainty. Hence, unexpected uncertainty can be used to estimate contextual uncertainty. Finally, we can note that while we didn’t explicitly write the effect of the distracting inputs for this non-stationary formulation, their contribution is also part of the Context bias^2^ term.

### 4 Explained and unexplained variance

Predictive models are commonly evaluated in terms of the fraction of variance unexplained or variation explained. Here we reiterate this formalism to provide perspective on the estimates of predictive uncertainty 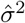 and confidence *c*, see equation (7). The expected squared prediction error *σ*^2^ between a prediction target *R* and a model prediction *Q* is a general metric of the prediction loss,

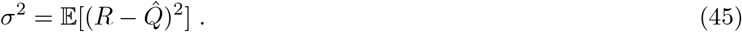

It is commonly also referred to as the residual variance or unexplained variance. This can be compared to the intrinsic variance of the target

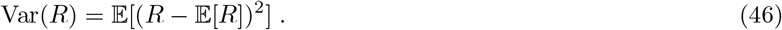

By taking the difference between the intrinsic variance and the unexplained variance, we get the variance explained VE (also referred to as explained variation)

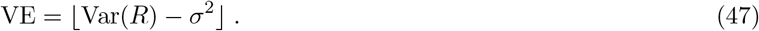

The VE quantifies how much the prediction error is reduced by the model 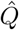 compared to using the unconditional expected value E[*R*] as an estimate. By dividing the unexplained variance *σ*^2^ with the intrinsic variance, we get the fraction of variance unexplained FVU,

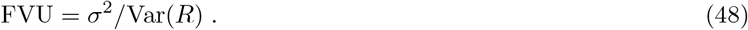

The analogous can be done to get the fraction of variance explained FVE, commonly referred to as coefficient of determination ℛ^2^,

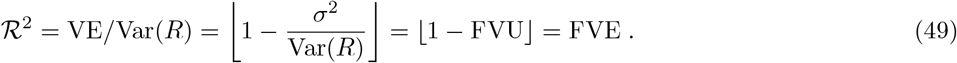

With this formalism at hand, the expected uncertainty 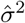, which is an online estimator of *σ*^2^, can be viewed as an internal representation of the unexplained variance. On the other hand, the prediction confidence

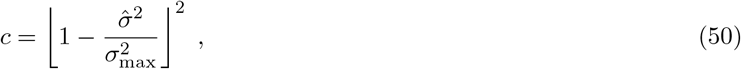

as defined in equation (7), represents an internal estimate of FVE^2^, the square of the fraction of variance explained, see equation (49). The rectifier function ensures that the confidence remains zero, even if the model 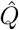 is a worse estimator than E[*R*]. While this situation would typically not be considered in stationary contexts, changes of context can lead the squared prediction error *σ*^2^ to rise above the intrinsic variance.

#### Note

Alternatively, we could have first introduced an objective prediction confidence as *c* = FVE^2^ and then the subjective estimated confidence as 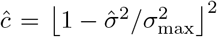. However, for simplicity we directly defined the internal representation of the prediction confidence as *c*, hence dropping the 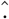 from *ĉ*, see equation (50).

### 5 Computing the reward variance

In the go/no-go sensory discrimination task, the distribution of the rewards *R* depend on the action selection probabilities. While it would be possible to continuously estimate the variance of the rewards for a changing distribution of action selection probabilities, this would require an online estimator. For simplicity, we use a fixed value 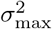, which was set to the variance in case of a uniformly random action selection policy *π*(*a*_GO_|*s*_1_) = *π*(*a*_GO_|*s*_2_) = *π*(*a*_NG_|*s*_1_) = *π*(*a*_NG_|*s*_2_) = 0.5. The value 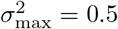 can be shown by

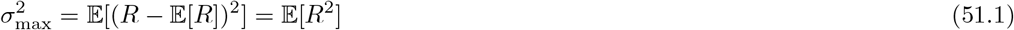

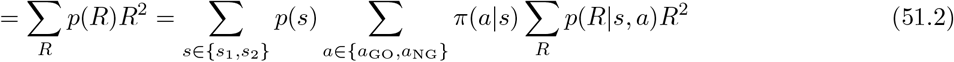

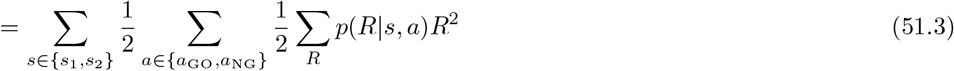

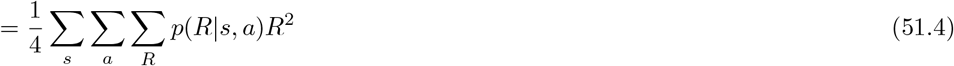

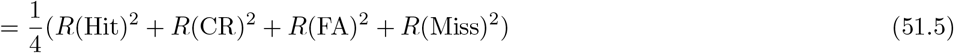

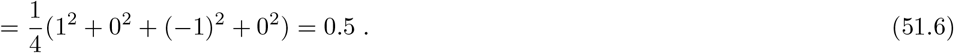

Equation (51.3) used the uniformity of stimulus sampling, meaning *p*(*s*_1_) = *p*(*s*_2_) = 0.5. Equations (51.5) and (51.6) used that, conditioned on action and stimulus, the reward outcomes are deterministic (*p*(*R*|*s, a*) ∈ {0, 1}), and the rewards *R* were {1, *™*1, 0} for Hit, FA and either CR or Miss respectively. Equation (51.1) used E[*R*] = 0. This can be demonstrated by

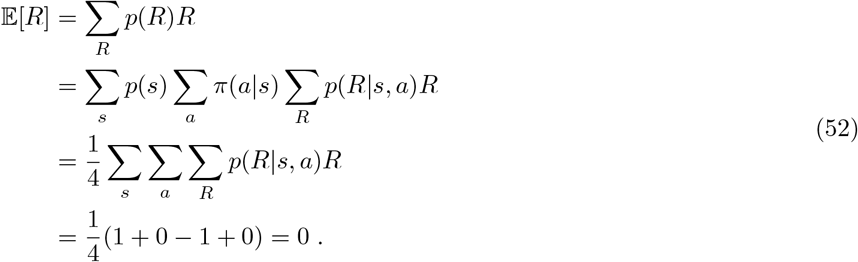

Note: in the same way that 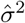 can be simultaneously interpreted both as the estimate of the variance unexplained by the predictor and as the expected prediction uncertainty, 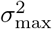 can be viewed under both lenses. On one side, it represents the variance of the reward, which can be used to compute the fraction of variance unexplained that appears in the coefficient of determination. On the other side, 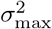 can be viewed as the prediction uncertainty of the most simple estimator 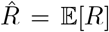. This explains, why the predictive confidence is zero, if the expected uncertainty 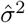 is greater than 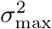, see equation (7).

### 6 Derivation of the confidence loss

The OFC activity at stimulus time is simulated as a prediction of the confidence loss CL. Confidence loss is modeled as being elicited by unexpected outcomes leading to a decrease in predictive confidence. The change in predictive confidence *c* is quantified by taking the difference in confidence between trial *t* + 1 and trial *t*:

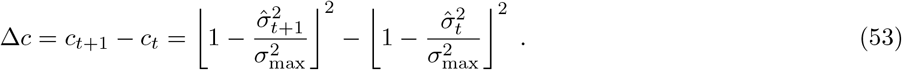

We ask how a change 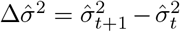 in the predictive uncertainty maps to a change Δ*c* in the predictive confidence. For this, we calculated the derivative of *c* with respect to 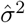,

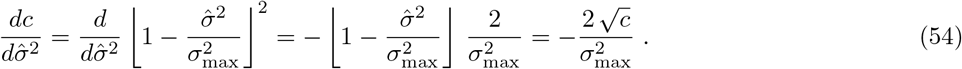

For small 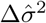 we can there for approximate

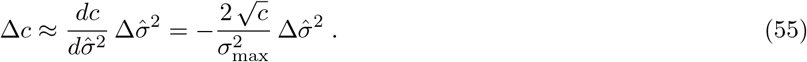

The same result can be obtained with finite time differences. For notational simplicity we assume that 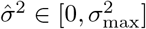, so that we can neglect the rectifier function ⌊·⌋. We then find

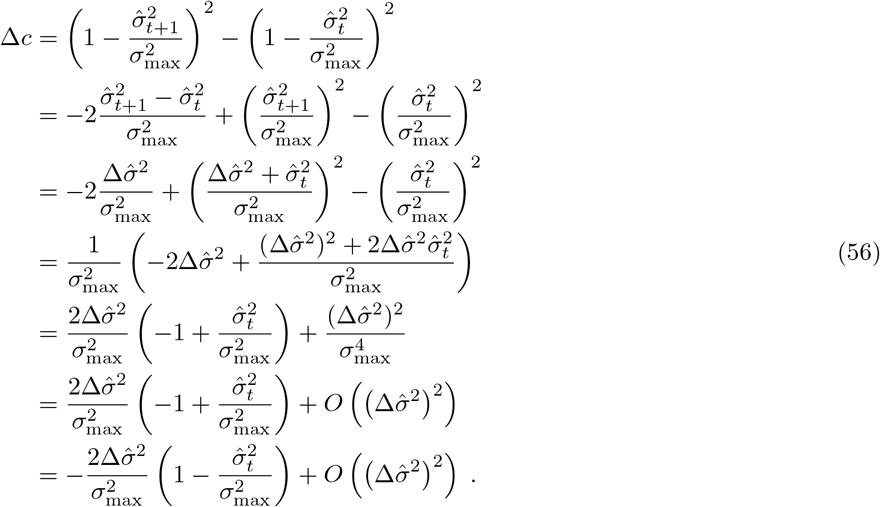

Confidence loss can therefore be approximated as

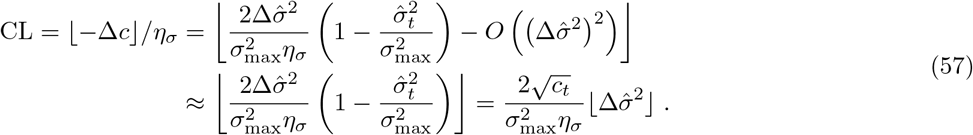

In the last step, we again made use of the fact that 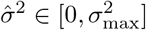 (or respectively that *c*_*t*_ ∈ [0, 1]).

